# Kinetic mechanisms for the sequence dependence of transcriptional errors

**DOI:** 10.1101/2025.03.04.641307

**Authors:** Tripti Midha, Anatoly B. Kolomeisky, Oleg A. Igoshin

## Abstract

The fidelity of template-dependent mRNA synthesis during transcription elongation is the primary determinant of accurate gene expression and the maintenance of functional RNA transcripts. However, the mechanisms governing transcription fidelity remain incompletely understood. While previous studies have characterized how error rates vary with nucleotide identity at upstream and downstream positions from the incorporation site, the comprehensive microscopic explanation of this sequence dependence has not been elucidated. In this study, we develop a novel theoretical approach that integrates transcription proofreading mechanisms and inhomogenous DNA sequence effects. Using first-passage analysis validated by Monte Carlo simulations, we quantitatively characterize nucleotide-specific error rates during RNA polymerase II transcription. The model accurately reproduces experimental error rates and predicts kinetic parameters influencing transcriptional fidelity. Analysis reveals nucleotide incorporation rates follow the hierarchy U*<*C*<*G*<*A, consistent with independent experimental observations. Notably, our model not only explains how the error rates depend on the nature of the base immediately down-stream (+1) but also predicts that the identity of the nucleotide at the second downstream position (+2) also plays an important role. Pyrimidines at position +2 contribute to lower error rates than purines, whereas the third downstream base (+3) has no effect. These previously unreported correlations are corroborated by bioinformatic analysis of existing datasets. In addition, using the BRCA1 gene as an example, we explore the physiological implications of sequence-dependent error rates, identifying an increased likelihood of premature stop codon errors. These findings clarify how DNA sequence context modulates nucleotide incorporation kinetics, advancing our understanding of transcriptional fidelity and its functional consequences.

## Introduction

The high fidelity of the transcription elongation process by RNA polymerase (RNAP) is crucial for the faithful gene expression (1). During transcription elongation, RNAP facilitates the addition of ribonucleotides (A, U, G, C) complementary to the DNA nucleotides (A, C, G, T) to the growing RNA chain by creating phosphodiester bonds. The process follows the Watson-Crick (WC) base pairing rules, implying that the purines (A, G) and pyrimidines (C, U) on the mRNA must make complementary base pairings with respective pyrimidines (T, C) and purines (G, A) on the DNA (see Fig. 1A) (2, 3). For accurate mRNA synthesis, the RNAP must distinguish the correct WC pairings (A:T, G:C, C:G, and U:A), denoted as the right, *R*, over the incorrect WC pairings, denoted as wrong, *W*, in the mRNA chain (see Fig. 1A) (3).

**Fig. 1.**
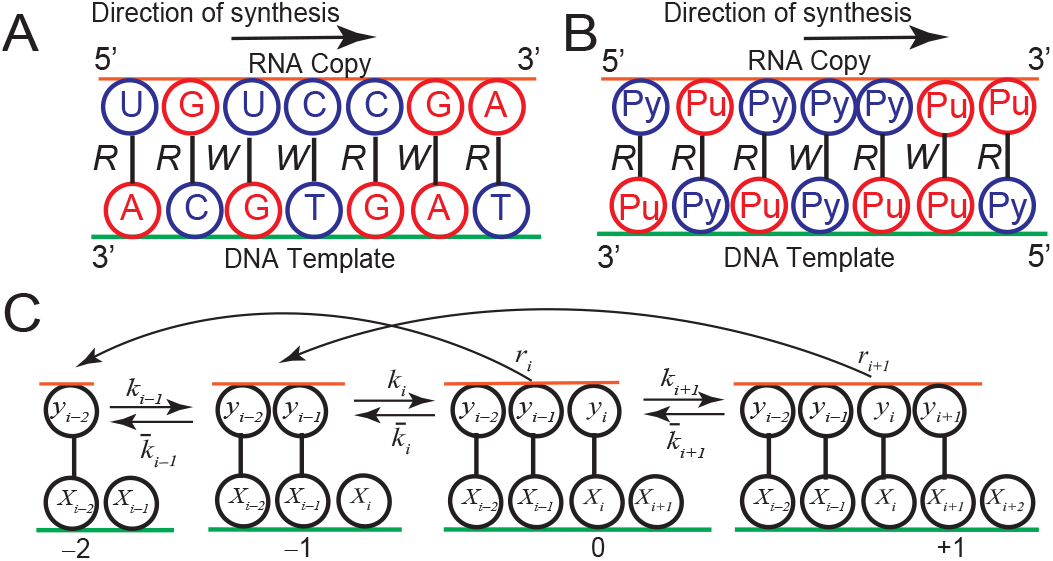
**(A)** (A) Watson-Crick (WC) base pairing rules between nucleotides on the DNA template and mRNA sequence during the transcription process with four nucleotides. **(B)** Watson-Crick (WC) base pairing rules in a transcription process with two components: purines (Pu), and pyrimidines (Py). **(C)** Schematic representation of the biophysical model for the transcription elongation process with effective forward, backward, and dinucleotide cleavage reactions.

Despite the inherent complexity of involved biochemical processes, RNAP achieves a remarkably low error rate, i.e., a fraction of incorrect pairings. In many biocatalytic processes, including transcription, the high accuracy arises due to out-of-equilibrium error-correcting mechanisms known as kinetic proofreading (KPR) (4, 5). KPR allows RNAP to kinetically discriminate between *R* and *W* substrates through proofreading steps (6, 7). RNAP possesses a spectrum of intrinsic proofreading tools, including pausing, backtracking, and endonucleolytic cleavage of dinucleotides, which do not rely on external energy sources (8–11). Together, these mechanisms enable to maintain high fidelity essential for cellular function (12–15).

Theoretically, the mechanisms of transcription elongation by RNAP have been well explored using different approaches such as Brownian ratchet, thermal ratchet (16, 17), and templated-copolymerization methods (7, 17–19). In the templated-copolymerization method, the elongation process is described using a homogenous Markov chain (18–22). Various other approaches, such as the Markov-chain copolymerization theory and linear-decoupling method, under steady-state conditions, have successfully predicted the fidelity and speed of the elongation process. However, all these methods considered the elongation process only as an *R/W* binary copolymerization process and, therefore, ignored the effects of nucleotide identity on the elongation and proofreading process. However, in reality, the DNA templates are heterogeneous, and each right and wrong pairing might have its own proofreading dynamics.

Recent experimental studies employing high-throughput techniques, such as circle sequencing, demonstrate that DNA template sequences strongly influence transcription error rates (23). While the overall trends are somewhat complex, it is clear that, compared to pyrimidines, purines at the next downstream position on mRNA (+1) increase the error rate at the current position (0) (23). Several recent studies on transcription elongation by the RNAP enzyme indicated that the incorporation rate of nucleotides follows the order U*<*C*<*G*<*A, suggesting that the incorporation rate of pyrimidines is slower than that of purines (16, 24). Furthermore, purines, double-ring structured nucleotides, have wider stacking areas than single-ring pyrimidines. This could increase their interaction strength with the active site of RNAP and influence their incorporation kinetics (25, 26). However, there is a lack of a theoretical understanding of the effect of nucleotide-specific kinetic rates on transcription fidelity for inhomogeneous DNA templates. Therefore, it is unclear if these kinetic differences can fully explain the sequence-dependence of error.

The sequence dependence of errors for inhomogeneous templates in DNA replication has been considered by several theoretical approaches, such as iterated functions and first-passage methods (27–30). These theories captured the potential role of nucleotide-specific kinetic parameters in controlling template-dependent fidelity (27–30). However, unlike DNA replication, transcription elongation involves more complex proofreading mechanisms involving backtracking and dinucleotide cleavage. Furthermore, there are fewer precise kinetic measurements for sequence-dependent transcription parameters. These reasons make theories for sequence-dependence of transcriptional errors more difficult to formulate.

In this paper, we develop a theoretical framework based on first-passage analysis that computes the template-specific transcription error rate and makes testable predictions of the kinetic parameters affecting transcription fidelity. First, in the context of the coarse-grained two-letter base alphabet (purines and pyrimidines), we examine whether our theoretical model can capture the experimentally observed sequence-dependent error rates (Fig. 2). We then use our theoretical approach to predict how the transcription fidelity is affected by long-range downstream neighbors, by extending the theory and checking the predictions by reanalyzing the data in Ref.(23) (Fig. 3). The results are utilized to predict sequence-dependent variation in kinetic parameters governing nucleotide incorporation and cleavage (Fig. 4). Furthermore, we also perform bioinformatics analysis to examine the physiological implications of sequence-dependent errors (Fig. 5). By examining the impact of synonymous errors and errors generating premature stop codons, our study provides insights into how transcription fidelity impacts functionality of living systems. Our study bridges theoretical understanding with experimental findings to unravel the complex mechanisms of transcription elongation.

**Fig. 2.**
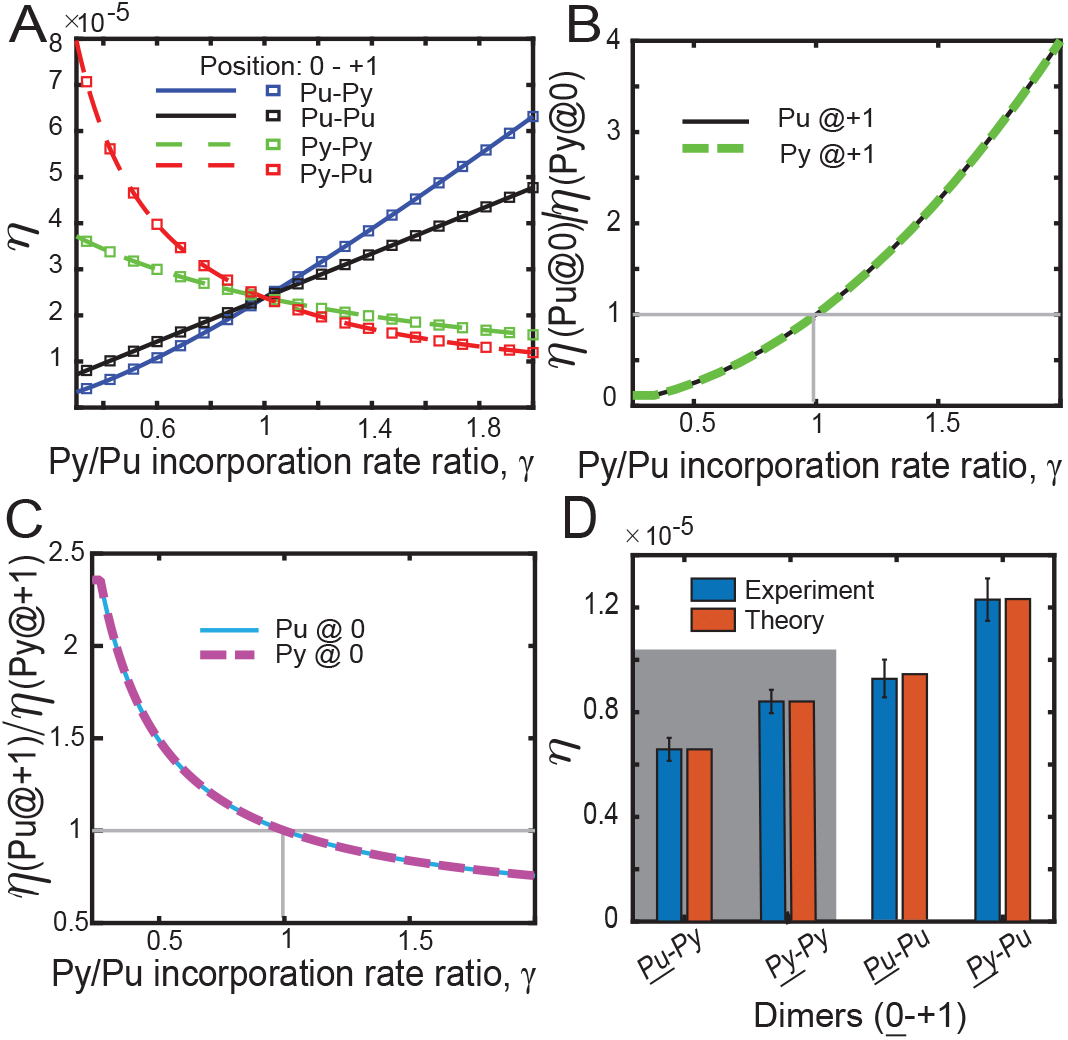
(A) Variation of error rate at position 0 as a function of the incorporation factor, *γ*, for four dimers occupying positions at 0 and +1 on mRNA for the model with two components. (B) Variation of the ratio of the error rates with purines at current position, 0, to the pyrimidines as a function of the incorporation factor, *γ*, for a fixed monomer at the next downstream position +1. (C) Ratio of the error rates with purines to the pyrimidines at the next downstream (+1) position as a function of the incorporation factor, *γ*, for a fixed monomer at the current position, 0. (D) Fitting (shaded region) of theoretical errors (red bars) from the model with first-order neighbor effects to the experimental data (blue bars). Theoretical error (red bars) at the optimal kinetic parameters for the unfitted (unshaded region) dimers aligns with the corresponding experimental data (blue bars). Error bars denote the standard error mean for the experimental data for the six samples (23).

**Fig. 3.**
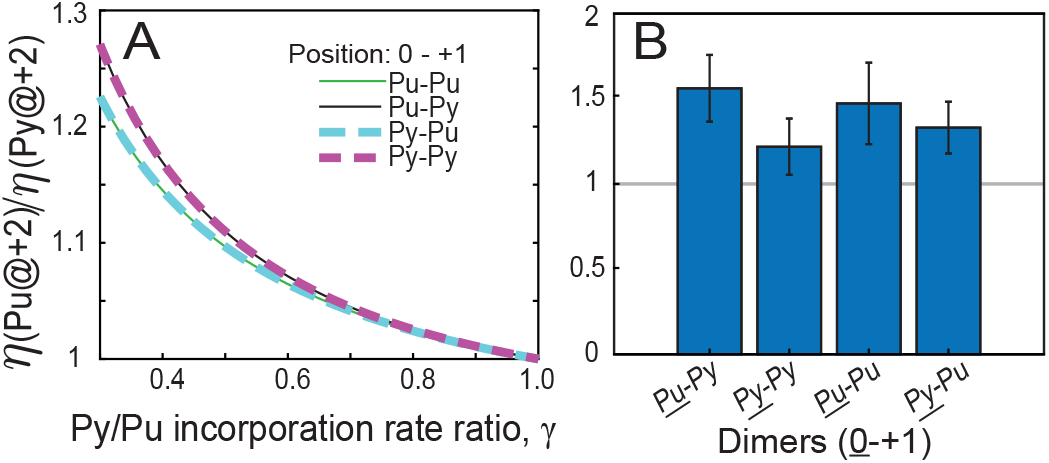
**(A)** Variation in the ratio of the theoretical error rates with Pu at +2 position to that of Py at +2 position for various dimers at 0, +1 position with changes in the incorporation factor, *γ*. **(B)** Ratio of the experimental error rates with Pu at +2 position to that of Py at +2 position for various dimers at 0, +1 position.

**Fig. 4.**
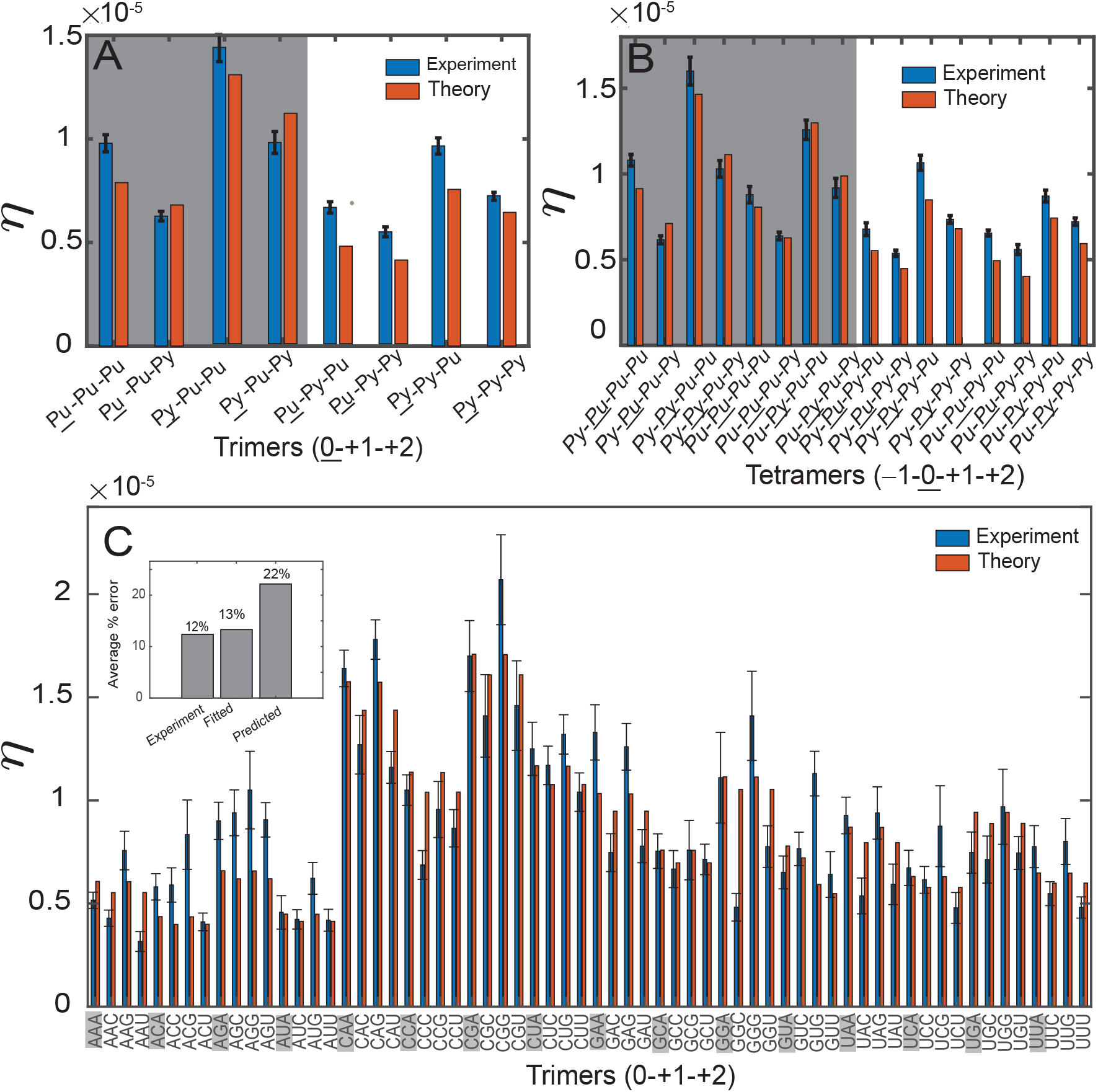
(A) Fitting of theoretical error rates 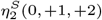 (red bars) depending on nucleotide identity at positions 0,+1,+2 to the corresponding experimental data (blue bars). (B) Fitting of theoretical error rates (red bars) 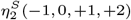 depending on the monomers at the -1, 0,+1,+2 positions to the corresponding experimental data (blue bars). (C) Fitting of theoretical error rates 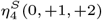 for the second-order biophysical model with four components to the corresponding experimental data. The inset shows the average error percentage for fitted, experimental, and unfitted data.

**Fig. 5.**
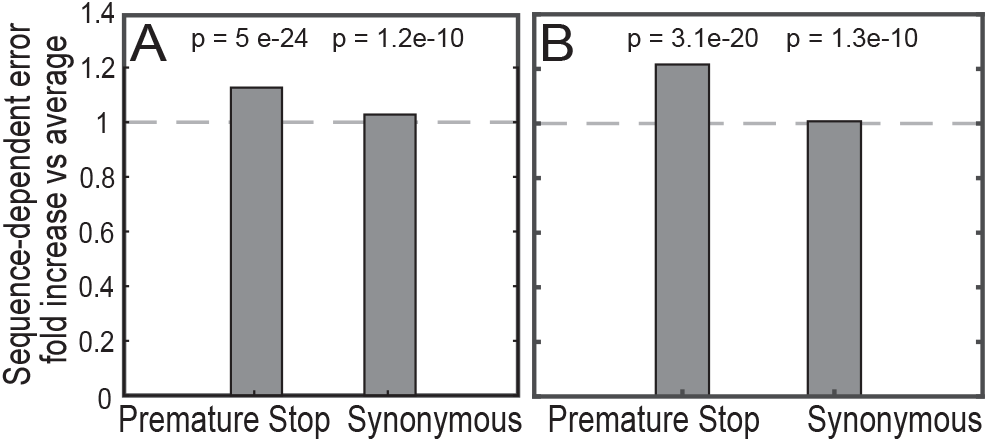
(A) Ratios of the error probability with experimental error depending on immediate downstream and upstream sequence to the error probability with non-sequence context dependent errors for the BRCA-1 gene. (B) Ratios of the error probability with experimental error depending on two immediate downstream neighbors and one upstream neighbor to the error probability with non-sequence context-dependent errors for the BRCA-1 gene.

## Results

### A. Nucleotide-specific kinetics and template heterogeneity modulate transcription elongation error rates

To analyze the effect of the chemical identity of the current base (0) and the chemical identity of the base immediately downstream (+1) on the error, we first employ an approximate biophysical model of transcription elongation (see Fig. S1, SI Appendix) with kinetic parameters as detailed in Table S1 (see Methods, SI Appendix). First, for simplicity, we aggregate the 4-letter base alphabet of nucleotides into the 2-letter description, i.e., we consider that both the DNA template and mRNA copy sequence consist of only two types of monomers: purines (Pu=A, G) and pyrimidines (Py =T, U/C). In that picture, it is also assumed that the correct (*R*) copying corresponds to associating Pu to Py, whereas the wrong (*W*) copying corresponds to associating Pu to Pu or Py to Py (see Fig. 1A, Fig.1B). It is important to note here that, as per the WC pairing rules, Pu:Pu and Py:Py bonds are always wrong, however, not all Pu:Py bonds are right (e.g. U:G). The limitation of the two-letter alphabet requires us to assume Pu:Py bonds are always right pairing, and this assumption allows us to pinpoint the intuitive basis of the sequence-dependence of errors. A more precise theoretical model with all four bases will be analyzed later.

The heterogeneity in the DNA template is introduced by assuming distinct incorporation rates of pyrimidines and purines, quantified by a non-dimensional scale-factor *γ*, i.e.,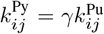. Here, the subscripts *i, j* ∈ {*R, W* } are two consecutive nucleotides on the mRNA chain, and the superscript denotes the nature of the incorporating nucleotide *j*, i.e, whether it is Pu or Py (see Methods for details). Using the first-passage analysis detailed in the Methods section, we numerically compute the template-dependent error rate, *η*, at the polymerase current position, 0. That error can be compared with the experimental data from Ref.(23) aggregated in the 2-letter description.

To predict how the difference in the nucleotide-specific incorporation rates affects the transcriptional error, we use the first-passage approach to compute the error rates for the four possible (Pu-Pu, Pu-Py, Py-Pu, Py-Py) nucleotide pairs (dimers) in the consecutive positions (0, +1) in the mRNA sequence. The results are shown in Fig. 2A as a function of *γ*. When *γ* = 1, i.e., for homogeneous kinetics, errors are the same for all dimers. For *γ >* 1, i.e., when the pyrimidines are incorporated faster than purines, the error *η* is highest for Pu-Py dimer and lowest for the Py-Pu dimer. Conversely, for *γ <* 1, *η* is highest for Py-Pu and lowest for the Pu-Py dimers. Monte Carlo simulations further confirm the accuracy of our predictions for heterogeneous DNA templates (Fig. 2A, lines vs symbols) (31). Notably, alternative approaches such as copolymerization(20) and linear decoupling(19) cannot correctly identify the error unless *γ* = 1, i.e., unless there is no sequence-dependence of polymerization rates. Thus, our theory predicts two effects: slower incorporation of a base at the current position increases the probability of error, whereas slower incorporation of a base at the next position decreases the error.

Notably, the effects of the current and the next bases on kinetics appear to be independent of each other. To explore the impact of the chemical nature of incorporating nucleotides at the current position, we compute the ratio of errors for dimers with Pu at position 0 to those with Py at position 0 (Fig. 2B). As one can see, this ratio remains the same regardless of the nucleotide at the +1 position. Similarly, to examine the influence of the nucleotide identity at the downstream position, we compute the ratio of errors for dimers with Pu at position +1 to those with Py at position +1 (Fig. 2C). This ratio does not depend on the identity of the nucleotide at position 0. Kinetic effects can be understood intuitively in the following way. For example, for *γ <* 1, the error rate is lower for purines than for pyrimidines in the current position because the incorporation of the wrong nucleotide (i.e. pyrimidine) is slower than that of the correct nucleotide (Fig. 2B). On the other hand, the error rate is lower for pyrimidines than the purines at the downstream base (Fig. 2C). After all, the slower bases at the downstream position give enough time to the polymerase to correct the error by cleaving nucleotides in case of misincorporations, thereby increasing the accuracy. Furthermore, the independence of the effects of the current and downstream bases (collapsing of the curves in Fig. 2B and Fig. 2C) can also be explained theoretically (see SI Appendix).

To quantitatively determine how the kinetic parameters for purines and pyrimidines differ, we fit theoretical predictions for the errors to corresponding experimental data. Note that the experimental data is known for the 64 cases, i.e., the error rates for nucleotides A, C, G, and U at a position (say) 0 for given nucleotides at the upstream (-1), and the downstream (+1) positions. We group them into four cases (Pu-Py, Py-Py, Pu-Pu, Py-Pu) occupying positions 0 and +1, as detailed in the SI Appendix. Moreover, to enable efficient parameter fitting, we utilize the approximate analytical expression of the theoretical error rate obtained under the biologically relevant conditions as a function of various kinetic parameters (see Methods, Eq. 7). This approximation (Eq. 7) agrees well with the results from the first-passage analysis (see Fig. S2). Given the sensitivity of transcription elongation errors to nucleotide incorporation and cleavage rates (7), we fix most of the parameters using previously determined values(7) as summarized in Table S1 and focus on fitting only three critical parameters: the incorporation rate for purines 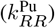, the pyrimidine-to-purine incorporation rate ratio (*γ*) such that 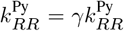, and pyrimidines to purines dinucleotide cleavage rates ratio, *ϕ*, such that *r*_Py_ = *ϕr*_Pu_. Here, *r*_*i*_ denotes the dinucleotide cleavage rates of misincorporated nucleotide when the correct one is *i* =Pu or Py. Using the least-squares optimization method weighted by the inverse of the standard deviation of experimental data, we fit theoretical error rates for Pu-Py and Py-Py dimers (shaded region in Fig. 2D) to experimental error rates by minimizing their differences. The results, displayed in Fig. 2D, show that the model successfully reproduces experimental error rates for dimers: Pu-Py, Py-Py, at the optimal parameter values: 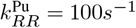, *γ* = 0.65, *ϕ* = 2.0, obtained from the fit. Based on the ensemble fitting of 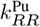, *γ* and *ϕ* (see SI Appendix), their coefficients of variations (CVs) are about 0.2, 0.1 and 0.2, respectively. Furthermore, error rates for the other two dimers not used in fitting, Pu-Pu and Py-Pu, are calculated using these parameters and fall within experimental uncertainty (unshaded region in Fig 2D). These findings indicate that pyrimidines are incorporated about 35% slower than purines (*γ*=0.65), but their cleavage rate is 2x higher. Note that a purely non-nucleotide-specific proofreading mechanism, i.e., when *ϕ* = 1, fails to fully account for the experimental data (see Fig. S3).

### B. Theoretical model and bioinformatics analysis reveal reduced transcription error with pyrimidines at the second downstream position on mRNA

During the proofreading process, RNAP has two possibilities to correct the error as it pauses twice – after the misincorporation of a nucleotide and after incorporating the following one. Each pause can result in dinucleotide cleavage, correcting the error (10). At each pause, there is a competition between the error correction and incorporation of the next base(7). Thus, sequence-dependence of incorporation rate couples the nucleotide identity at the second downstream position (+2) and the error rate. To predict these effects theoretically, we generalize our theory to account for second-order effects, i.e., we now consider the dependence of kinetic rates on the presence of errors in the two preceding bases and the nature of the incorporated bases (SI Appendix, Fig. S4). The error rate for the more advanced model can also be computed numerically using the first-passage approach or with an approximate analytical formula (see Eq. 8, and SI Appendix).

To quantify the influence of the second downstream nucleotide, we compute the ratios of the errors with Pu at +2 position, *η*(Pu@ + 2), to that of Py at +2 position, *η*(Py@ + 2), for all four possible dimers at preceding positions. Fig. 3A shows the results as a function of *γ*. We observed that for *γ <* 1 (as indicated in the previous section), these error ratios consistently exceed 1. Thus, the error rate is higher when purines are at the +2 position because the slower incorporation of pyrimidines at +2 allows more time for error correction. This effect is independent of the nucleotide identity at position 0 but is influenced by the identity at position +1. Specifically, this error ratio is greater when Py occupies position +1 than Pu. Consequently, the error is lowest for the Py-Py combination at positions +1 and +2 (SI Appendix, Fig. S5).

It is important to note that while the authors of Ref. (23) did not report the effect of the +2 position on error, their accumulated data allows us to quantify these effects and test our theoretical predictions. To this end, we conducted a bioinformatics analysis of experimental data from Ref. (23) (see Fig. S6A). In Fig. 3B, we plot the ratio of the experimental error rates with Pu at +2 position to those with Py at +2 position for all four possible dimers at positions (0, +1). Across all dimers, the ratio exceeds 1, indicating that pyrimidines at the second downstream position reduce the error as predicted by our theoretical results. Thus, we demonstrate the influence of the second downstream base pair on the error.

Since RNAP cleaves only dinucleotides, it is expected that the identity of the nucleotide at the third downstream (+3) position from the site of incorporation (0) should not significantly impact the error rate. To test this hypothesis, we compute the ratio of the error with purines at +3 downstream position to the error with pyrimidines at the same location using Eq. 8. Indeed, the ratio is 1 if the identities of the nucleotides at the preceding positions are the same (see Fig. S7A). To validate these theoretical results, we again performed bioinformatics analysis of the experimental errors as a function of the identity of the nucleotide at position +3. The results show that the ratio of the errors with purines and pyrimidines at +3 position is always 1 within the error bar range (see Fig. S7B). These results indicate that the transcriptional error rate has correlations only up to the second base, reflecting the processes utilized in proofreading (namely, dinucleotide cleavage).

### C. Predictions of the long-range genetic-context dependent errors align with experimental data

Next, we asked if our more advanced theoretical model could explain all corresponding experimental data and predict which kinetic parameters are sequence-dependent. For this purpose, we fit the theoretical errors from Eq. (8) for four (Pu-Pu-Pu, Pu-Pu-Py, Py-Pu-Pu, Py-Pu-Pu) of the eight trimers occupying positions (0,+1,+2) to the corresponding experimental data. Note that *η* in Eq. 8 is the function of various kinetic parameters. However, we choose to optimize the same kinetic parameters as in Fig. 2 fit, i.e. the incorporation rate for purine 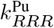, *γ* such that 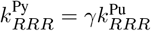, and the dinucleotide cleavage rate of misincorporation instead of correct Pu, denoted by, *r*_Pu_. Here, in 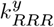, the superscript *y ∈* {Pu, Py} denotes the nature of the incorporating nucleotide, whereas the subscript denotes a trimer in the mRNA chain. The remaining parameters are fixed as described in Table S2.

The results in Fig. 4A indicate that for the optimal kinetic parameters, the theoretical errors are in excellent agreement with the corresponding experimental data for all the fitted trimers: Pu-Pu-Pu, Pu-Pu-Py, Py-Pu-Pu, Py-Pu-Pu (shaded region in Fig. 4A) as well as for the remaining predicted trimers: Pu-Py-Pu, Pu-Py-Py, Py-Py-Pu, Py-Py-Pu (unshaded region in Fig. 4A). Furthermore, the estimate for the incorporation rate parameter 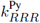 is found to be close to the experiments and the previously discussed first-order model (10).

While our model naturally captures the effect of the downstream neighbors on the error rate, the identity of the upstream neighbor doesn’t directly affect the error (see Fig. S6B). This result contrasts with our bioinformatics analyses of the experimental data of Ref. (23) (see Fig. S6A). This is because, in our model, the incorporation rate of any nucleotide at the current position (0) depends on whether the nucleotide at the upstream position (-1) is right or wrong, but not on its chemical identity. Thus, to account for the observed effects, the identity of the upstream base pair should also affect the incorporation rate. However, how the upstream base affects the incorporation rate is not known. Therefore, we choose the simplest implementation, in which the incorporation rates of any nucleotide at position 0 are multiplied by the factor *ψ* if its upstream (-1) neighbor is a pyrimidine. The resulting approximate expression for the second-order model with two components, *η*(− 1, 0, +1, +2) (Eq. 8), will also depend on parameter *ψ*. To estimate the parameter *ψ*, we use the data on error rate depending on the identity of four positions (−1, 0, +1, +2) and fit half of these values to our theoretical predictions (shaded region of Fig. 4B) using the non-linear weighted least square method. At the optimal value of *ψ* ≈ 0.88, the fitted theoretical error (shaded region) and predicted theoretical error (unshaded region of Fig. 4B) are in reasonable agreement with the experimental data. Thus, we predict a slight 12% deceleration of the incorporation rate when the upstream (-1) nucleotide is a pyrimidine.

We next evaluate if our model generalized to a 4-letter description, i.e., with four monomers (A, T, C, G) on the DNA template binding with the corresponding nucleotides (U, A, G, C) on the mRNA sequence (see Fig. 1A), can explain the corresponding experimental data and predict the incorporation rates of individual nucleotides. For this purpose, we computed the error rates under the biologically relevant condition (see SI Appendix), as given in Eq. (9). The theoretical error rate is a function of various parameters (given in Table S3). We choose to find the optimal value of the 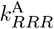 for nucleotide *A*, and the ratios *γ*_1_, *γ*_2_, *γ*_3_ describing relative incorporation rates for other nucleotides, such that 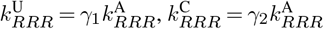, and 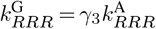. We also find the optimal fit for the cleavage rates of nucleotides in place of correct A, C, G, and U, denoted by *r*_*A*_, *r*_*C*_, *r*_*G*_, and *r*_*U*_, respectively. Using the non-linear weighted least square approximation method, we fit the theoretical error for analytical expression of error rate under the biologically relevant conditions for every fourth trimer in the 64 combinations (shown by the shaded trimers in Fig. 4C) as given in Eq. 9, to the corresponding experimental data in Fig. 4C. The optimal values for the parameters obtained from the fitting are given in Table S4. The robustness and uniqueness of the inferred parameters from the fitting were also confirmed with the ensemble fitting; CVs from the resulting distribution are shown in Table S4. At these optimal values, the prediction for the trimers not used for fitting (unshaded trimers in Fig. 4C) also agrees well with the experimental data, with a mean relative percentage error of ∼22%. The fitting results indicate that the incorporation rates of nucleotides are in the following order: U*<*C*<*G*<*A, which also agrees well with the experimental findings (16, 24). Moreover, our fitting analysis reveals that the cleavage rate for misincorporations replacing correct pyrimidines (C) is higher than that for misincorporations replacing correct purines (G), aligning with the previously discussed two-letter alphabet model. Note that, for the fitting, we assume that the cleavage rates depend on the nature of the correct nucleotide being replaced by any misincorporated nucleotide. The fitting results for the case when cleavage rates depend on the nature of the misincorporated nucleotide are shown in Fig. S8, SI Appendix.

### D. Sequence-dependent error increases the likelihood of premature terminating codon errors

To investigate the physiological significance of the sequence-dependent error rate, we asked how these effects might influence proteins after translation. As an example, we decided to focus on the 22 coding exons (coding sequences (CDS)) of the BRCA-1 gene, transcribed by RNAP. The human BRCA-1 gene has several isoforms encoding proteins of different lengths. For instance, the BRCA1-201 encodes the full-length protein consisting of 1863 amino acids and plays a central role in DNA repair and tumor suppression. Transcriptional errors can have distinct effects on the proteins (32, 33). On the one hand, it can change a codon to that encoding for the same amino acid (e.g. UCU (serine) changes to UCC (serine)), resulting in synonymous errors; on the other hand, it can change it to a stop codon (e.g., CAA (Glutamine) changes to UAA (a stop codon)), resulting in premature termination. We first investigated the physiological implications of the sequence-dependent errors for different nucleotides at positions -1, 0, and +1. For this purpose, we examined the error probability, *E*, defined as the fraction of the BRCA1 transcripts affected by premature stop errors or synonymous errors. Assuming that the error rates for each nucleotide are independent, for each value of the error rate, we computed the sequence-dependent error probability *E*_*gc*_(−1, 0, +1) = 1 − Π_*j*_(1 − *n*_*ijk*_*η*_*ijk*_), of the occurrence of at least one error in the gene. Here, the product runs over index *j*, representing each nucleotide in the sequence that leads to any error, *i* denotes the 5’ (upstream) neighbor of *j*, and *k* denotes the 3’ downstream neighbor of *j*. Parameters *n*_*ijk*_ and *η*_*ijk*_ denote the number of errors and the error rate of the nucleotide *j*, respectively, in the sequence (*i, j, k*). We compare this error probability with a weighted average error rate for the BRCA-1 gene, given by *E*_*avg*_ = 1 − Π_*j*_(1 −*n*_*j*_*η*_*avg*_(−1, 0, 1)). Here, *η*_*avg*_(−1, 0, 1) is computed as the weighted average of the sequence-dependent error rate (*η*(−1, 0, +1)) of all nucleotides in the genome. We then found the normalized error probability, *E*_*avg*_, of all isoforms of the BRCA1 gene by dividing the error probability by the length of the amino acid sequences. Figure 5A displays the ratio of the mean of these normalized error probabilities, 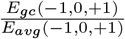 for both premature stop errors and synonymous errors, where *η*_*ijk*_ is taken from the experimental data. We found the ratio to be always greater than 1, but the effect is much stronger for the premature stop errors. It is also determined that this observation is statistically significant by computing their p-values, which are small as indicated (see Fig. 5A). A similar behavior on the ratio of the error probabilities is observed when the long-range genetic context (-1,0,+1,+2) dependent error rates are considered (Fig. 5B). These results indicate that the sequence-dependent error, whether short-range or long-range, increases the likelihood of premature termination in the encoded protein.

## Discussion

In this work, we developed a theoretical framework to quantify the error rate of RNAP by explicitly considering the sequence dependence of transcription kinetics. We first considered a simplified model with first-order neighbor effects with only two types of components: purines and pyrimidines, on both the mRNA sequence and the DNA template (Fig. 1). The sequence-dependent error rate was computed using the first-passage method, and it aligns well with the Monte Carlo simulations (Fig. 2). Further fitting of theoretical predictions against experimental data shows a good quantitative agreement, supporting the model’s ability to capture the sequence-dependent errors observed in transcription (Fig. 2).

The analysis suggests that the incorporation rates of pyrimidines are slower than purines, whereas the cleavage rate of misincorporated purines is faster. To account for the effect of second-downstream (+2) neighbors on the error rate at the current position (0), we performed the analysis for a more advanced (second-order) biophysical model (Fig. 3). Our results demonstrated that the error rate at each site is significantly impacted by the chemical identity of the immediate downstream and the second downstream bases. Our findings reveal that pyrimidines at downstream positions enhance transcriptional accuracy due to their slower incorporation rates and associated proofreading processes. In contrast, the identity of the third downstream base does not have any impact. These predictions are validated by performing bioinformatics analyses of the published experimental data. When extended from a two-component to the four components (A, C, G, U), our models also accurately predict nucleotide-specific incorporation and proofreading kinetics (see Table S4), as validated by the experimental data (16, 23, 24).

Furthermore, with bioinformatic analysis, using BRCA1 coding sequences as an example, we demonstrated that sequence-dependent errors might significantly increase error probabilities for premature stop errors. These findings underscore the functional implications of transcription fidelity and highlight the necessity of integrating sequence-dependent effects into transcription models to accurately predict broader physiological consequences.

Our theoretical approach to investigate transcription elongation, calibrated against experimental data, accurately predicts nucleotide-specific incorporation rates. The analysis reveals that purines (A, G) polymerize faster than pyrimidines (C, U), a trend supported by experimental findings (25, 26). Structurally, purines have a double-ringed configuration, which facilitates stronger base-stacking interactions, probably leading to faster rates of association compared to single-ringed pyrimidines, where such interactions are weaker (25, 26). Experimental studies further suggest that polymerization rates correlate with the 3’ terminal base composition, following the order U*<*C*<*G*<*A. Additionally, a sequence-dependent thermal ratchet model for transcription elongation supports this hierarchy, reinforcing that pyrimidine incorporates more slowly than purines (16, 24). Our advanced theoretical model with second-order neighbor effects and all four nucleotides predicts the same incorporation rate hierarchy (U*<*C*<*G*<*A) as in the experiment. To further validate and refine the model, future experimental studies, such as sequence-specific cleavage rate assays, could target specific perturbations to incorporation or cleavage pathways, enabling independent measurement of individual rate constants.

Our theoretical analysis of errors under biologically relevant conditions suggests that, in addition to nucleotide incorporation rates, the dinucleotide cleavage rate of misincorporated nucleotides must be sequence-dependent. This cleavage is the basis of proofreading and mostly occurs after incorporation. By fitting our theoretical model to the corresponding experimental data, we found that dinucleotide cleavage rate for misincorporations replacing correct pyrimidines is approximately twice as fast as the cleavage rate for those replacing correct purines. This value agrees with experimental studies showing that errors occurring in place of pyrimidine (U) are removed at twice the rate of those replacing purine (G) (12). Furthermore, the dinucleotide cleavage mechanism introduces two proofreading checkpoints that enhance transcriptional accuracy (7). This suggests that transcriptional fidelity is influenced not only by the identity of the nucleotide at the immediate downstream position but also by the identity of the second downstream nucleotide (see Fig. 3). Our more advanced theoretical model with second-order neighbor effects reveals that error rates decrease when pyrimidines occupy the second downstream position and reach their lowest when two consecutive pyrimidines are present. Conversely, the highest error rates occur when two purines appear in succession. This is consistent with the fact that pyrimidines exhibit slower incorporation rates and higher proofreading efficiency compared to purines (12, 24). Thus, transcriptional accuracy is highest when two pyrimidines are positioned consecutively, whereas error rates peak when two purines appear together.

Experimental studies indicate that the incorporation rates of nucleotides are affected when the downstream neighbor to be incorporated is a pyrimidine (34). However, it is unknown to what extent the incorporation dynamics will be affected when pyrimidines are at the upstream position. The experimental data emphasize the role of both upstream and downstream neighbors in determining transcriptional error (23); hence, the upstream neighbor effects on the incorporation dynamics must be determined. Our results indicate that incorporation rates of nucleotides are reduced by a factor *ψ* when the upstream (-1) nucleotide is a pyrimidine. In particular, our advanced model accurately reproduces experimental observations at the optimal *ψ ≈* 0.88. This value suggests that pyrimidines at upstream positions slightly slow down the incorporation process, thus potentially enhancing proofreading capability and transcription fidelity.

While transcriptional errors were the main focus of the study, our formalism can also assess the effects of nucleotide-dependent rates on transcriptional speed. Our results show that with increased fractions of purines, both mean first-passage time (MFPT, i.e., the inverse sequence-dependent transcription speed) and its coefficient of variation (CV_*T*_) increase (Fig. S9). This effect may be biologically important, as genes involved in rapid stress responses may benefit not only from fast average transcription but also from reduced timing variability (35, 36).

Transcriptional errors, influenced by genetic context, can affect the error probability (37–40). We examined the physiological implications of sequence-dependent error rates by assessing their impact on error probabilities in the BRCA1 gene, a key tumor suppressor (32, 33). Premature stop codon errors can impair protein function (32), while synonymous errors may reduce translation efficiency (33). Our analysis revealed that sequence-dependent error rates result in higher error probabilities than non-sequence-specific errors. This suggests that the context sequence plays a significant role in determining transcription errors. The effect is mild for synonymous errors but about 20% for premature termination errors. The higher error probability for premature termination can be attributed to the composition of stop codons UAA, UAG, and UGA, which all contain a pyrimidine at position 0 and purines at the +1 and +2 downstream positions. Since purines at these downstream positions are associated with the highest error rates, there is a higher chance of misincorporations changing other bases into U at position 0, resulting in a stop codon, i.e., premature termination errors. These findings highlight the relationship between transcription fidelity and functional genomic outcomes, stressing the critical role of genetic context in biological information processing. In conclusion, this study bridges theoretical insights with experimental data, providing a foundation for predictive models of transcription fidelity. Our results collectively emphasize the importance of integrating both short- and long-range correlations and sequence-specific kinetic parameters in models of transcription fidelity.

In this study, we have modeled dinucleotide cleavage during transcription proofreading as an irreversible process to keep the kinetic scheme analytically tractable. While this simplification allows us to quantify fidelity through first-passage kinetics, it omits the potential thermodynamic cost of reversible proofreading cycles. In future work, extending the model to include reversibility would, for each nucleotide, allow us to quantify the trade-offs between error and the excess dissipation in each proofreading cycle, i.e., extra entropy production per incorporation step (41). Such a treatment could connect transcription fidelity with the fundamental principles of nonequilibrium stochastic thermodynamics (42, 43).

## Methods

We consider a theoretical framework for a template-dependent transcription elongation process, in which the monomers are added to the growing end of the mRNA molecule according to the DNA template sequence. We denote the sequence of length *L* on the DNA template as *X*_1_*X*_2_ … *X*_*L*_. The elements of the mRNA copy sequence are denoted by *y*_*i*_ for *i* = {1, 2, … *L*}. During elongation, each DNA base *X*_*i*_ pairs with an mRNA base *y*_*i*_, forming *X*_*i*_ : *y*_*i*_ pairs. The accuracy of these pairings is characterized by a binary sequence *α*_*i*_ where *α*_*i ∈*_{*R, W*} indicates whether the pairing at position *i* is correct (*R*) or incorrect (*W*) (see Table S5 for the symbols definition).

We first consider a biophysical model incorporating first-order neighbor effects, with both the DNA and mRNA sequences composed of two nucleotide classes: purines (Pu) and pyrimidines (Py),i.e., *X*_*i*_, *y*_*i*_ ∈ {Pu, Py}. In this model, the kinetic rates for nucleotide addition (*k*_*i*_), removal 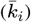, and dinucleotide cleavage (*r*_*i*_) at position *i*, depend on the accuracy of both current and the previously incorporated nucleotide, *α*_*i*_ *α*_*i −*1_, i.e., whether *α*_*i*_, *α*_*i −*1_ are *R* or *W* (see Fig. S1A). Additionally, we assume that the rates *k*_*i*_ and *r*_*i*_ also depend on the chemical identity of the incorporating nucleotide *y*_*i*_, i.e., Pu or Py (see Fig. 1C, Fig. S1B). It should be noted that if *α*_*i*_ = *R*, and *X*_*i*_ = Pu, then the corresponding incorporated nucleotide *y*_*i*_ = Py, if *α*_*i*_ = *R*, and *X*_*i*_ = Py, then *y*_*i*_ = Pu, *α*_*i*_ = *W*, and *X*_*i*_ = Pu, then *y*_*i*_ = Pu, and if *α*_*i*_ = *W*, and *X*_*i*_ = Py, then *y*_*i*_ = Py. In our model, we consider the incorporation rates of pyrimidines and purines to be related by 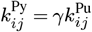, where, the superscript reflects the nature of incorporating monomer *y*_*j*_, i.e., Pu or Py.

Our theoretical framework for transcription elongation is based on mean first-passage time analysis (29, 35, 36), wherein we compute the sequence-dependent transcription error rates by analyzing the stochastic dynamics of RNA polymerase as it transcribes a heterogeneous DNA template. We describe transcription elongation as a first-order Markov chain process from a reflecting boundary at the first position to the absorbing boundary at the last. This description allows us to view the transcription elongation as a first-passage process. We assume that the mRNA transcription proceeds with the preexisting initial seed *R* or *W*, i.e., either correctly paired with the 3’ end of the DNA template or not. The chain grows as the monomer at position *i* is added and removed with the rates *k*_*i*_ and 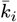, respectively. However, as boundary conditions, the chain’s initial seed and the last added monomer cannot be removed. Additionally, to account for proofreading, we consider dinucleotide cleavage with the rate *r*_*i*_ from all sites except *i* = 1, 2, and *L*. In our model, the kinetic rates are the effective rates obtained by combining several reaction substeps (3, 7). We now write the temporal evolutions of probability, 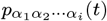, of the growing chain sequence *α*_1_ *α*_2_ … *α*_*i*_ (1 *≤ i ≤ L*) for the given DNA template *X*_1_*X*_2_, … *X*_*L*_, at time *t*, for the bulk positions *i* (3 *≤ i ≤ L* − 3),

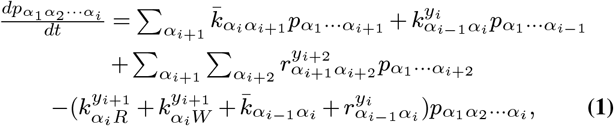

Here, the summations runs for *α*_*i*+1_, *α*_*i*+2_ ∈ *R, W* . The temporal evolution equations for the boundary positions are provided in the SI Appendix. We are interested in finding the final sequence distribution of the nascent chain, i.e., the long-time limit 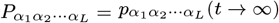. For this, the following boundary conditions are assumed. The initial boundary condition is 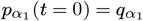, such that *q*_*R*_ + *q*_*W*_ = 1 for a given *X*_1_ at position 1 on the DNA template, 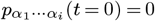 (for *i ≥* 2). We also assume the transcription will be completed fully to absorbing state *L* in the long time limit, i.e., 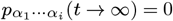 for 1 ≤ *i < L*. One can then integrate Eq. 1, along with the temporal evolution equations for the probabilities for the boundary positions (see the SI Appendix) for *t* varying from limit 0 to ∞, and this leads to the iteration relations as given in SI Appendix. They can be transformed into an intuitive form of a forward inhomogeneous Markov chain, as below

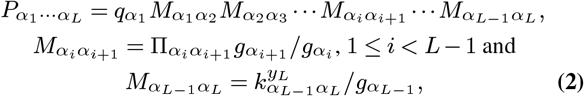

The expressions for 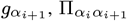 are provided in SI appendix. Furthermore, following the approach of Ref. (29), we can introduce the 2 × 2 stochastic transfer matrix **M**_*i,i*+1_ with each row sum equal to 1, i.e., 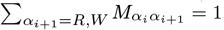, for *α*_*i*_ = *R, W* . In this context, one can calculate the positional probability, 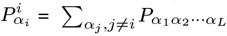, for *α*_*i*_ = *R* or *W* at any position, *i*, on mRNA sequence given the monomer *X*_*i*_ at position *i* on the DNA template by 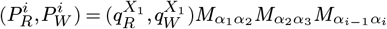. For the first-order process, with one right and wrong nucleotide for a given monomer at the DNA template, one can write **M**_*i*−1,*i*_, as

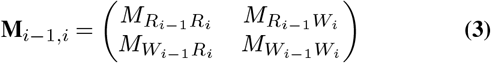

Equivalently, the probability distribution at site *i* is given as, 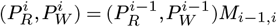 (30). In particular, one can obtain the error probability,

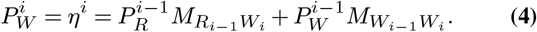

Our approach also allows us to calculate the mean first-passage time (MFPT) for the RNAP to reach the end of the chain. It is defined as

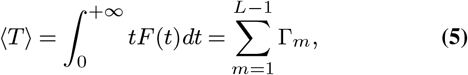

where *F* (*t*) is the first-passage time distribution function to reach the position *L* for the first time starting from the position 1, determined by the equation

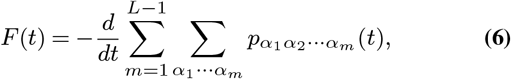

and 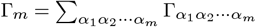 (see SI Appendix for details). The analytical results for MFPT are validated using stochastic simulations, which also provide the full first-passage time distribution. From these, we compute the coefficient of variation (CV_*T*_) to quantify temporal variability in elongation timing (see SI Appendix, Fig. S9).

We now obtain the approximate analytical expression for the error rate under the biologically relevant conditions inspired by the kinetic parameters of real RNAPs (3, 8, 12). The analytical expressions of error are utilized to fit the theoretical results to the experimental data.

For the biophysical model with first-order neighbor effects and two components (Fig. S1), the biologically relevant conditions are intuitive as follows:

1. 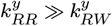, implying the addition of *R* nucleotide is always much faster than that of *W*, irrespective of the identity *y* of the incorporating monomer.
2. 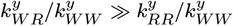, implying that the successive additions of *W* are almost inhibited, taken to approximately zero.
3. 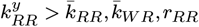. This means that successive additions of *R* always dominate the backward and cleavage rate, irrespective of the identity *y* of the incorporating monomer.
4. 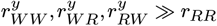, which means the terminus containing *W* s are likely to be proofread faster.

Under the biologically relevant conditions, we first approximate 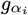 ‘s. The condition 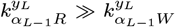 implies 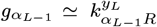. Following the same logic, we obtain 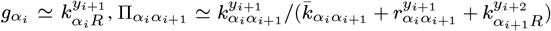. Similarly, one can write the approximate expression for the individual elements of the matrix *M*_*i*_ (see the SI Appendix). Further, under the biologically relevant conditions, *M*_*RR*_ ≫ *M*_*RW*_, *≫ M*_*WR*_ ≫ *M*_*WW*_, and *M*_*RW*_ ≫ *M*_*WW*_ (19, 29, 30). The analytical expression of the probability for a wrong nucleotide at any *i*^*th*^ position of the chain is thus given as 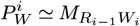 (see Appendix C, Ref. (29)). For the first-order biophysical model with two components, we denote the approximate expression of error rate at any current site, 0 (say), depending on the nucleotide at the downstream position, +1, by 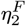, where the superscript represents the first-order model and subscript represent the two components. That is,

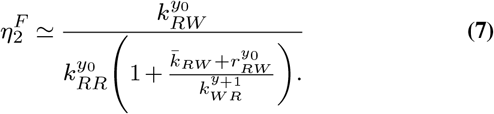

Here, the denominator of the RHS reflects the dinucleotide cleavage and the first-order neighbor effects. Depending on the chemical identity of the nucleotides *y*_0_, *y*_1_ ∈{Pu, Py}, we obtain the theoretical error rate for four dimers (*y*_0_-*y*_+1_) (see SI Appendix).

The theoretical error rates were fitted to experimental measurements to infer kinetic parameters governing transcriptional fidelity. Parameter optimization was performed using MATLAB’s derivative-free patternsearch algorithm, which minimizes the weighted sum of squared residuals between theoretical and experimental error rates. Residuals were scaled by the corresponding experimental standard deviations to account for measurement uncertainty (see SI Appendix for details). To assess robustness and potential non-uniqueness of parameter inference, we performed ensemble fitting with 1000 randomly initialized optimization runs, sampling starting values within biologically plausible bounds. The ensemble fitting revealed a sharp peak in the parameter distribution, suggesting that the inferred values are well-constrained and uniquely determined by the experimental data.

For the more advanced model with second-order neighbor effects and two components (Pu, Py), the rates *k*_*i*_, 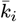, and *r*_*i*_ depend on the accuracy of 3 nucleotides, i.e. on *α*_*i−*2_, *α*_*i−*1_, *α*_*i*_ (see Fig. S4A). For the template-dependent kinetics, we consider that the rate for addition, *k*_*i*_, also depends on the identity of the incorporating nucleotide *y*_*i*_ = Pu, Py. Thus, the incorporation rate 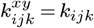, if *y* = Pu, and 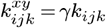, if *y* = Py. Here, *i, j, k* = *R, W*, *x* = Pu or Py, is the identity of the previous nucleotide *j*, and *y* denotes the nature, i.e., Pu or Py, of the current incorporating nucleotide *k*. Additionally, we also consider the case when the identity of the preceding (upstream) nucleotide *y*_*i*−1_ = Pu or Py can also affect the incorporation rate *k*_*i*_ (SI Appendix, Fig. S4B). For this case, when upstream nucleotide identity effects are considered, the rate 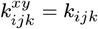 if *x, y* = Pu, 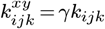 if *y* = Py and *x* = Pu, 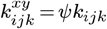 if *y* = Pu and *x* = Py, 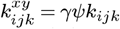 if *y* = Py and *x* = Py. We start the elongation process with the initial seed as dimer *α*_1_*α*_2_ with probability 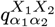, such that 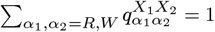. Following the same logic as for the first-order biophysical model, we can obtain the final sequence probability 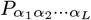 for the second-order model as a forward inhomogeneous Markov chain (see SI Appendix for details). Under the biologically relevant condition (see the SI Appendix), the approximate expression of the error rate for the second-order biophysical model with two components, 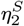, where the superscript represents the second-order model and subscript represents the two components, can be written as,

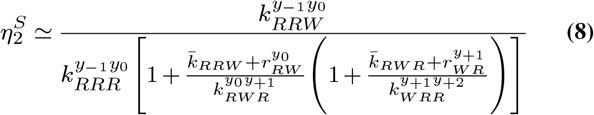

Here, the denominator in the RHS reflects the dinucleotide cleavage and the second-order neighbor effects on the error rate. For the case with no upstream nucleotide identity effects, we calculate the error rate for 8 trimers (*y*_0_−*y*_+1_−*y*_+2_) depending on the chemical identity of the nucleotides *y*_*i*_ ∈ {Pu, Py} for *i* = {0, +1, +2} (see Fig. 4A). For the case with upstream nucleotide identity effects, we calculate the for 16 tetramers (*y*_−1_ −*y*_0_ −*y*_+1_ −*y*_+2_) depending on the chemical identity of the nucleotides *y*_*i*_ ∈ {Pu, Py} for *i* = {−1, 0, +1, +2} (see Fig. 4B).

To consider a more realistic case when both DNA and mRNA sequences consist of four components (A, C, G, T/U), for every correct pairing *R* (A:U, C:G, G:C, T:A) we need to account for the three wrong pairing possibilities *W*_*j*_, for *j* = 1 to 3. The kinetic parameters for the addition and the removal of every correct nucleotide and their corresponding three wrong nucleotides for the second-order biophysical model are given in Table S3. Similarly to the second-order model with two components, one can obtain the error for the model with four elements. Here, we write the approximate analytical expression of the error rate for the second-order model with four components, 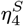, where the superscript represents the second-order model and the subscript denotes the four components, under bio-relevant conditions (see the SI Appendix) as below.

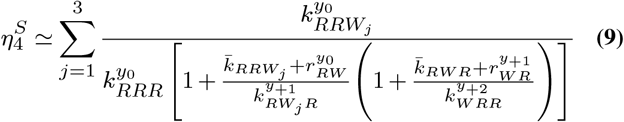

Using the above equation, one can obtain the error rate for the 64 trimers (*y*_0_− *y*_+1_− *y*_+2_) depending on the chemical identity of the nucleotides *y*_*i*_ ∈ {A, U, C, G} for *i* = {0, +1, +2} (see Fig. 4C). We sincerely thank the authors of Ref. (23) for generously providing the datasets from their study. This work was supported by the Center for Theoretical Biological Physics National Science Foundation (NSF) Grant PHY-2019745. OAI also acknowledges support from the Welch Foundation Grant C-1995. ABK also acknowledges the support from the Welch Foundation (C-1559).

## Supporting Information Text

### Biophysical model with first-order neighbor effects

Here, we write the temporal evolutions of probability, 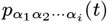, of the growing chain sequence *α*_1_ *α*_2_ … *α*_*i*_ for *i* = 1, 2, *L* − 2, *L* − 1, *L* as below

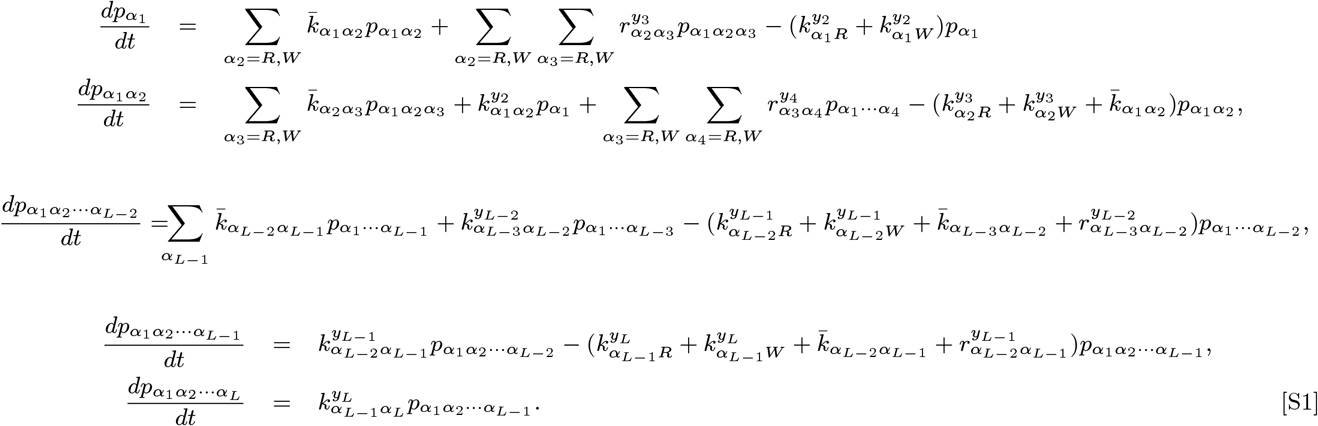

Here, we obtain the final sequence distribution, i.e., 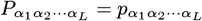 (*t → ∞*) of the mRNA chain. For this purpose, we integrate the temporal evolution equations for 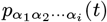, ∀*i* = 1 to *L*, given by Eq. (S1), and Eq. 2 (main manuscript), under the boundary conditions as defined in the methods (main manuscript).

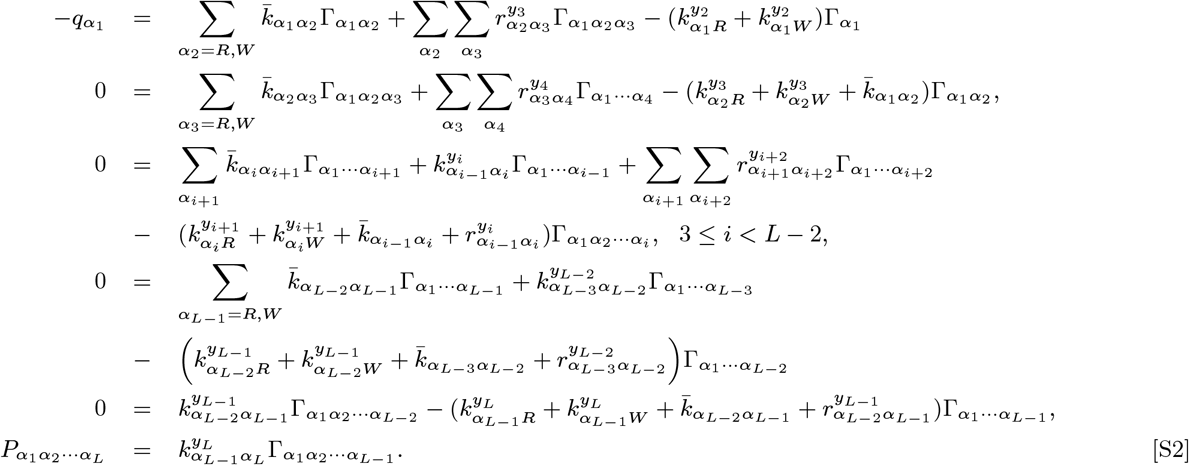

Here,

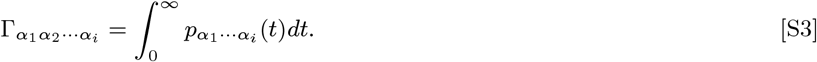

We now solve the above equations to obtain 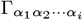 as

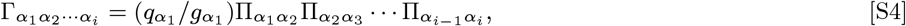

and the following iteration relations:

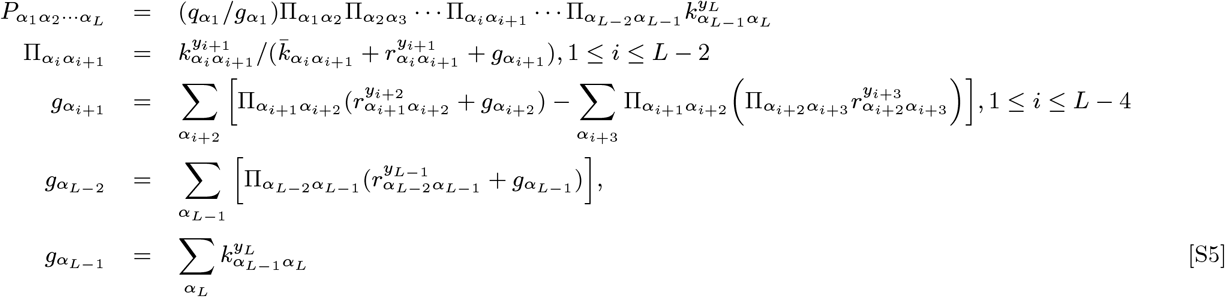

The above iteration relations are further utilized to compute the stochastic transfer matrix **M**_*i*−1,*i*_. For instance, under the bio-relevant conditions (see methods), we can obtain 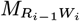, which is nothing but an approximate probability of a wrong nucleotide 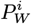 at position *i*, and is given as

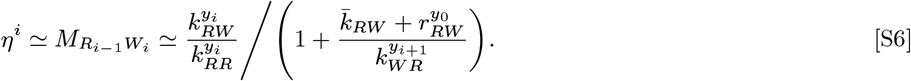

### Mean first-passage time and coefficient of variation

We consider the transcription elongation process as a first-passage process from the reflecting boundary at a starting site *α*_1_ to the absorbing boundary at the last site *α*_*L*_. For this first-passage process, we calculate the mean first-passage time (MFPT) to reach the end of the mRNA chain of length *L* using our theoretical approach. The MFPT is defined as the expected value of the first-passage time (FPT), *T*, and is thus written as

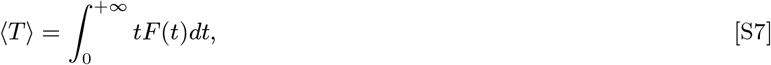

where *F* (*t*) is the first-passage time distribution from position 1 to *L*. One can express first-passage time in terms of the survival probability, *S*(*t*), given by the relation 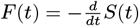 (1). Here, *S*(*t*) is defined as the total probability of the system in any non-absorbing state at time *t*, and is given as

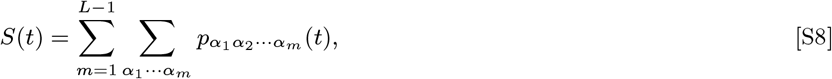

where 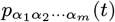 is the probability of the growing chain *α*_1_*α*_2_ … *α*_*m*_ at position *m*. We now apply integration by parts to Eq. S7, and further utilize the boundary conditions as defined in the main manuscript, Eq. S8, and Eq. S3, to obtain

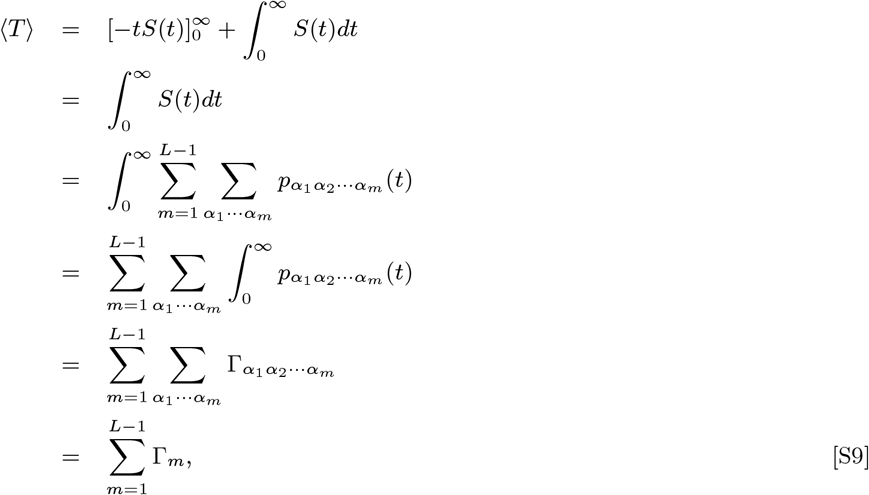

where 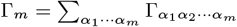 . Using Eqs. S4 and Eq. (2) (main manuscript), one can write

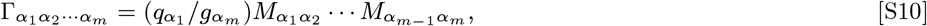

In another form, Γ_*m*_ can be written as

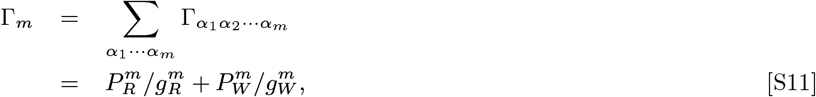

where 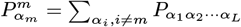, denotes the positional probability for *α*_*m*_ = {*R, W* } at position *m* on the mRNA chain.

To quantify the stochastic variability in transcription timing beyond average elongation speed, we computed the coefficient of variation (CV_*T*_) of the first-passage time (FPT) distribution for each sequence context defined as

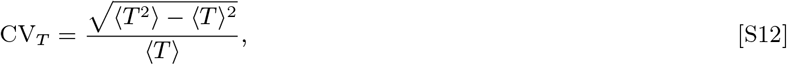

where both ⟨*T* ⟩, and ⟨*T* ^2^⟩ are estimated from stochastic simulations. While we derived analytical expressions for the MFPT, calculating higher-order moments such as ⟨*T* ^2^⟩ is analytically intractable for sequence-dependent dynamics and complex proofreading pathways. Instead, we simulated full transcription trajectories using the Gillespie algorithm and extracted the FPT distributions directly (see SI Appendix, Monte Carlo simulations section). This allowed us to compute CV_*T*_ values for various DNA templates, revealing that purine-rich sequences tend to exhibit greater timing variability compared to pyrimidine-rich or mixed templates (Fig. S9). These results suggest that sequence composition not only shapes fidelity and speed but also modulates the consistency of transcription timing.

### Parameter inference and ensemble modeling

To estimate kinetic parameters governing error rates in our theoretical model, we employed the patternsearch optimization method. For a given set of parameters *θ* to be predicted, we define the residual vector *Res* as the weighted difference between the theoretically predicted error rates and the corresponding experimental data as given below

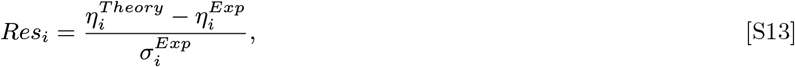

where 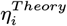 is the error rate obtained from the theory, 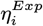 is the corresponding experimental data, and 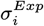 is the standard deviation of the experimental value. The objective function minimised is the sum of squared residuals, and is given as

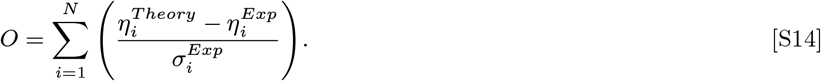

This is equivalent to the weighted nonlinear least squares method, incorporating experimental uncertainty as weights. To evaluate the robustness and potential uniqueness of the inferred parameters, we performed ensemble optimization with 1000 randomized initial conditions. We also quantified the dispersion of each parameter using its coefficient of variation, defined as CV = Standard deviation*/*Mean. A low CV indicates that the optimization consistently converges to similar parameter values, suggesting uniqueness and identifiability.

### Theoretical explanation for the identical error rate ratio for dimers in the first-order biophysical model with two elements

In this section, we mathematically justify the reason for the two identical error ratio curves in Fig. 2B, and Fig. 2C (main manuscript) for the first-order biophysical model with two elements. For this purpose, we first theoretically compute the error ratio: *η*(Pu@0)*/η*(Py@0) (see Fig. 2B, main manuscript) and show that it is identical for either Pu at +1 position or Py at +1 position. We then compute the error ratio *η*(Pu@ + 1)*/η*(Py@ + 1) (see Fig. 2C, main manuscript) and show that the ratio is identical for either Pu at 0 position or Py at 0 position. To show this, we write the analytical expression of error rate for dimers *y*_0_ −*y*_+1_, occupying positions (0, +1), where *y*_0_, *y*_1_ ∈ {Pu, Py} for the first-order biophysical model (see methods, main manuscript),

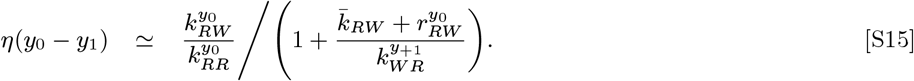

Explicitly, the error rate, *η*(*y*_0_ − *y*_1_), for all possible dimers *y*_0_ − *y*_1_ at (0,+1) positions can be written using Eq. S15 as below

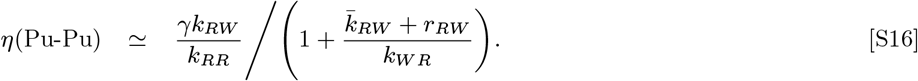

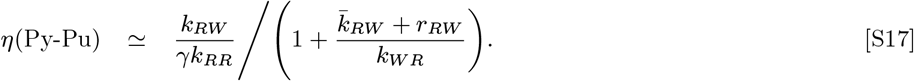

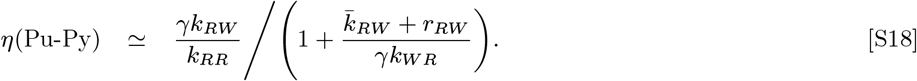

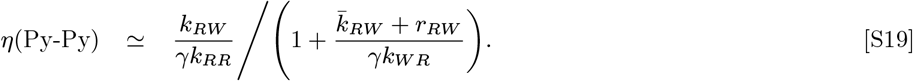

Using the above equations, we obtain

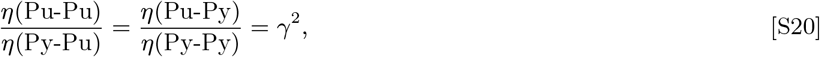

for non-nucleotide-specific dinucleotide cleavage rates. Whereas, in the case when the cleavage rate for purines and pyrimidines differ by a factor *ϕ*, we get

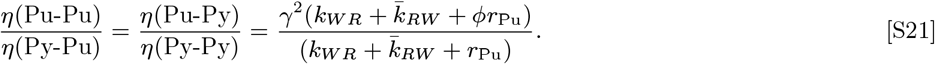

It is clear from Eqs. (S20)-(S21) that the ratio of the error rate with Pu @0 to Py @0 is the same whether there is a Pu or Py at the +1 position. This is the reason we get identical curves, as shown in Fig. 2B (main manuscript) for the case of non-nucleotide-specific proofreading. Similarly, we observe that

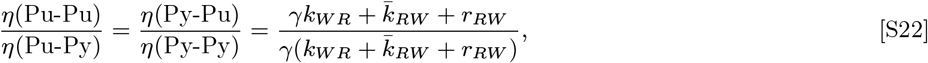

for non-nucleotide-specific dinucleotide cleavage rates. This implies that the ratio of the error rate with Pu @+1 to Py @+1 is the same whether there is a Pu or Py at the 0 position. This is the reason for the identical two curves in Fig. 2C for the case of non-nucleotide-specific proofreading. Thus, it can be said that the effects of current-base and next-base kinetics are independent of each other.

### Advanced model with second-order neighbor effects

For the theoretical model with second-order neighbor effects, we assume that the elongation process starts with the initial seed *α*_1_*α*_2_, with probability 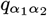, such that 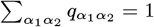. The first-passage process description for the model assumes that the first dimer acts as a reflecting boundary, i.e., it can not be removed, and the last site is the absorbing boundary. Let *α*_1_*α*_2_ *α*_*i*_ denote the growing *mRNA*, with probability 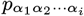 for a given DNA template *X*_1_*X*_2_ & *X*_*i*_. We write the temporal evolution equation for these probabilities, as below,

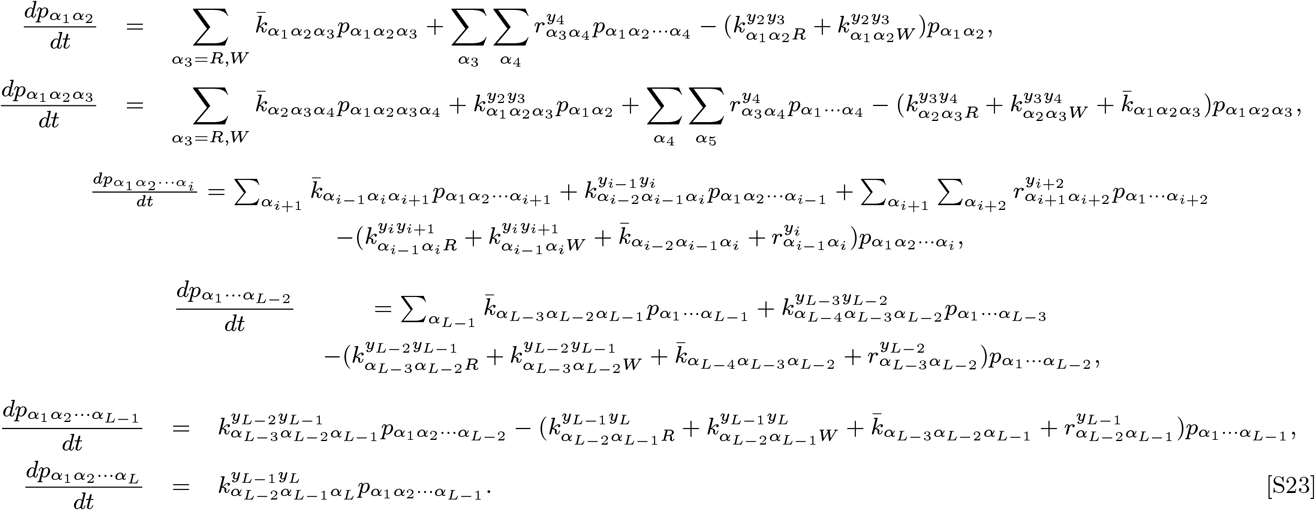

We now obtain the final sequence distribution, i.e., 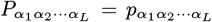 (*t→ ∞*) of the mRNA chain for the second-order model, under the following boundary conditions. The initial boundary condition is 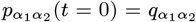, such that 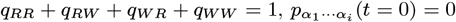 (for *i ≥*3). We also assume the long time limits 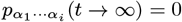 for 2 ≤ *i < L*. We next integrate the temporal evolutions equations for 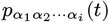, ∀*i* = 1 to *L*, given by set of Eqs. (S23), to obtain

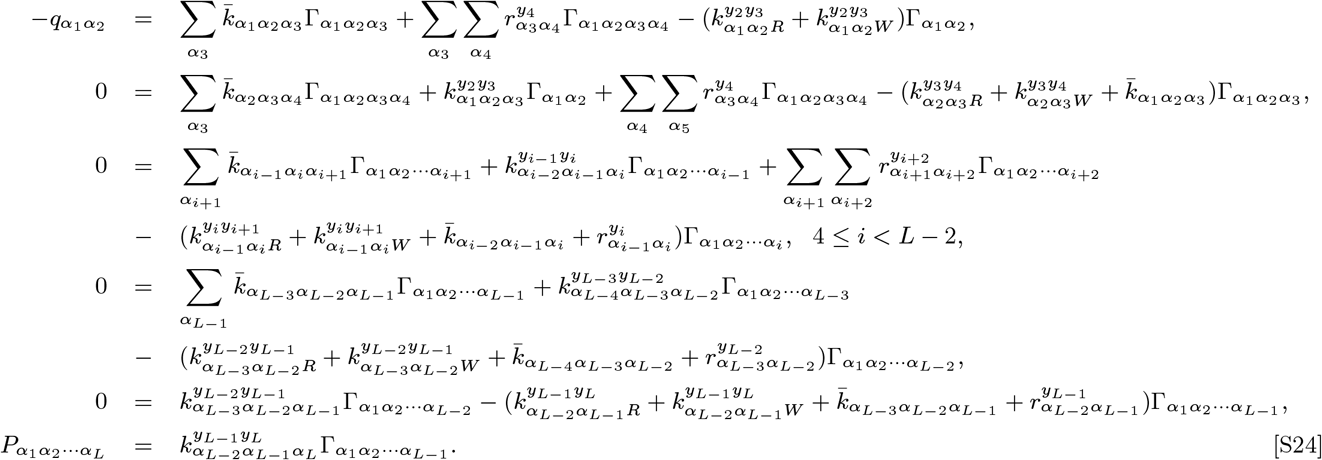

Here,

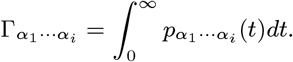

One can solve the above equations to obtain the following iteration relations:

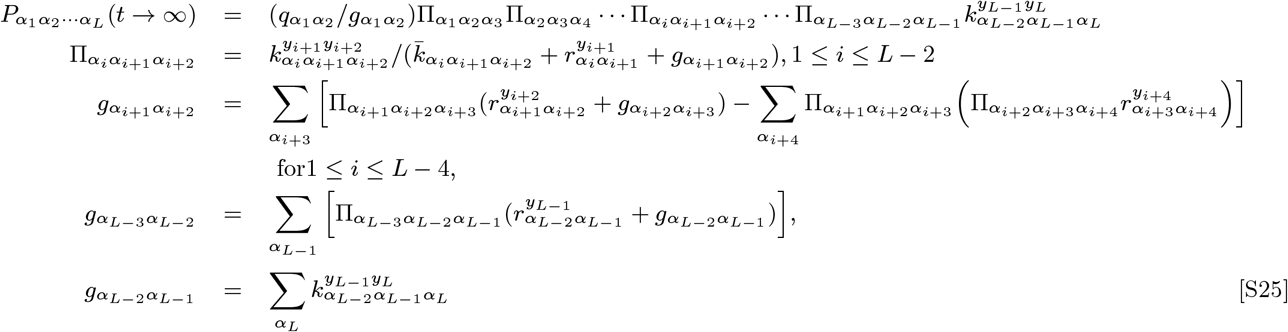

Equivalently, the above equations can be transformed into an intuitive form of a forward inhomogeneous Markov chain, as below

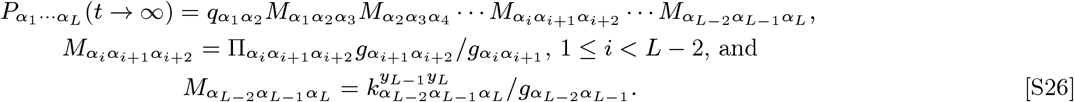

Following the approach in Refs. (1, 2), one can obtain the second-order stochastic transfer matrix **M**_*i*−2*i*−1*i*_

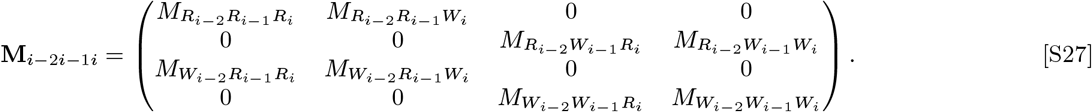

Next, to obtain the approximate expression of the elements of the above matrix, we define the biologically relevant conditions for the advanced model with second-order neighbor effects as intuitive below (1, 3):

1. 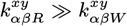, implying the addition of *R* nucleotide is always much faster than that of *W*, where *α, β* = *R, W*, and *x, y* = Pu, Py.
2. 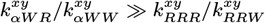, implying that the successive additions of *W* are almost inhibited, taken to approximately zero. In fact, 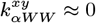.
3. 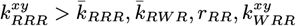. This means that successive additions of *R* always dominate the backward and cleavage rates, guaranteeing the transcriptional elongation.
4. 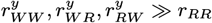, which means the terminus containing *W* s are likely to be proofread faster.

Note that for the model with four components, where we have 3 possible *W*_*i*_’s for every R incorporated, the above state bio-relevant condition for every *W*_*i*_, for *i* = 1, 2, 3. Under the above-stated biological relevant conditions, one can calculate 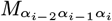 by the following iterations 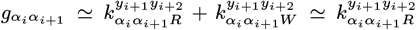, and 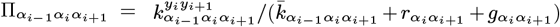. Also we get 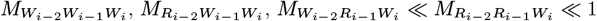. In this approximation, the positional probability 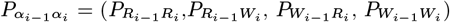 represented by the eigen-vector, 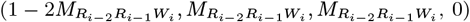, of the matrix *M*_*i*−2*i*−1*i*_ can be obtained (2). Equivalently, the probability of a wrong nucleotide at position *i, η*, is given as 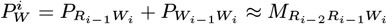, where

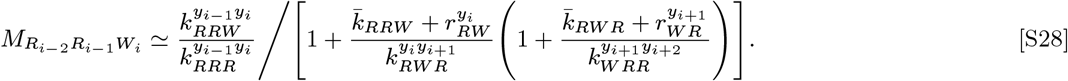

Thus, the approximate analytical expression of the error for the advanced model with second-order neighbor effects is

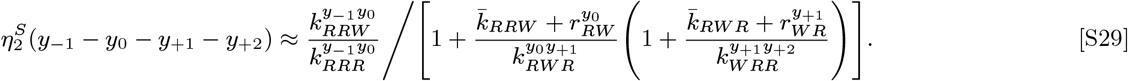

### Grouping of experimental data for 64 cases based on nucleotides at -1,0,1 positions into 4 cases

In this section, we describe how the experimental error rates from Ref. (4) for nucleotides A, C, G, and U at position 0 were systematically grouped based on the identity of the nucleotides at the upstream (-1) and downstream (+1) positions. The goal of this procedure is to categorize the error rates into four distinct cases, depending on whether the nucleotide at position 0 is a purine (A, G) or a pyrimidine (C, U), and whether the nucleotide at position +1 is a purine (Pu) or pyrimidine (Py). We first collected the error rates for each nucleotide *i* at position 0, considering every possible nucleotide *j* at position +1, while disregarding the identity of the nucleotide at position -1, for *i, j* ∈{A, C, G, U}. This resulted in 16 distinct cases, accounting for all combinations of the four nucleotides at positions 0 and +1. Next, we grouped the data based on whether the nucleotide at position +1 was a purine (A, G) or a pyrimidine (C, U). For each nucleotide *i* at position 0, we separately considered its error rates when the downstream nucleotide was either a purine or a pyrimidine, reducing the number of cases from 16 to 8. Finally, we categorized the data based on the type of nucleotide at position 0. This resulted in 4 cases: Pu-Pu, Pu-Py, Py-Pu, Py-Py occupying 0, +1 positions on the mRNA sequence.

### Bioinformatics analysis of the experimental data for error rate depending on the bases at one upstream and two downstream positions

We group the experimental error rates from Ref. (4) for nucleotides A, C, G, and U at position 0 based on the identity of the nucleotides at the upstream (-1), first downstream (+1), and second downstream (+2) positions. This grouping will categorize the error rates for 256 cases arising from nucleotide identity at (-1,0,+1,+2) positions into 16 distinct cases. These cases arise depending on whether the nucleotides at the positions (-1,0,+1,+2) are purine (A, G) or pyrimidines (C, U). We first collected the error rates for each nucleotide at position 0, considering whether the nucleotide at position -1 is a purine or a pyrimidine, reducing the cases to 128. We then grouped the data based on whether the nucleotide at +1 is purine or pyrimidine and then based on whether the nucleotide at +2 position is purine or pyrimidine. Finally, we categorized the data based on the type of nucleotide at position 0, resulting in error rates for 16 cases depending on whether there is a purine or a pyrimidine at -1, 0, +1, and +2.

### Monte Carlo Simulations

To complement the numerical results for our theoretical model for two elements, purine (Pu) and pyrimidine (Py), we perform extensive Monte Carlo simulations (MCS). We employed the Gillespie algorithm (5) to simulate the growth of mRNA chain for a given DNA template sequence of length 1000. The mRNA sequence for the first-order neighbor effects starts with a monomer as an initial seed and grows according to the kinetic rates given in Table S1. For the process with second-order neighbor effects, the mRNA sequence is extended from a dimer as an initial seed. The probability of errors is directly inferred from the outcomes of 1000 simulations. A length of 1000 monomers is chosen to avoid any edge effects in computing the probability of errors.

We also used Gillespie stochastic simulation to model the kinetic trajectory of RNA polymerase transcribing a DNA template of length *L* = 1000. For each trajectory, the total time taken to reach the terminal state (position = *L*) was recorded as the first-passage time (FPT). We ran 10^5^ independent simulations and compiled the distribution of FPTs. From these, we calculated the mean first-passage time (MFPT), variance, and coefficient of variation (CV_*T*_).

---

**Fig. S1.**
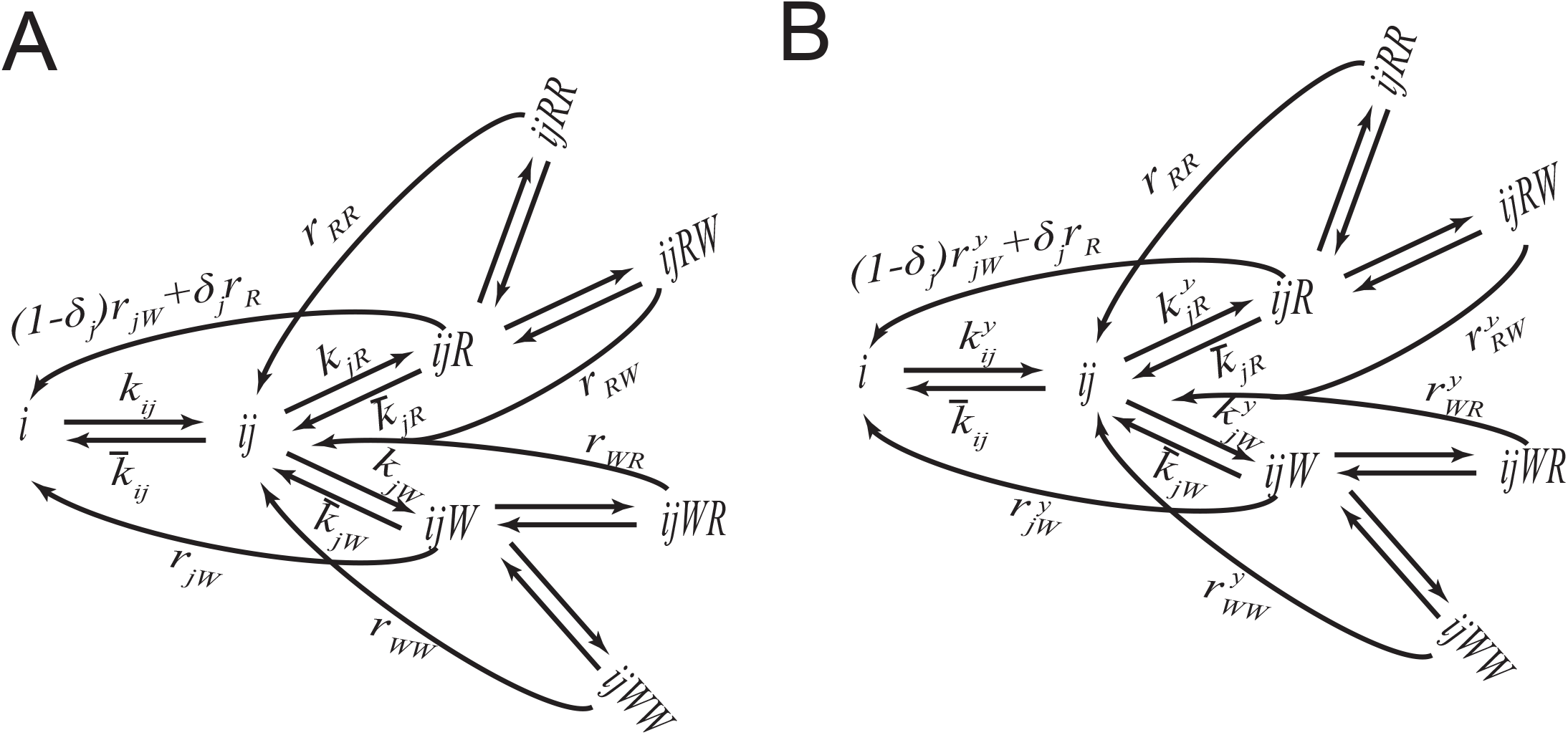
Schematic diagram of the two-elements first-order biophysical model for transcription elongation process with dinucleotide cleavage and (A) without template-dependent kinetics (B) with template-dependent kinetics. Here, *i, j* = *R, W*, *i* denotes the initial monomer in the mRNA sequence, and *y* denotes the nature, i.e., Pu or Py of the current incorporating nucleotide *j*. The rate 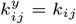, if *y* = Pu, and 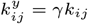, if *y* = Py.

**Fig. S2.**
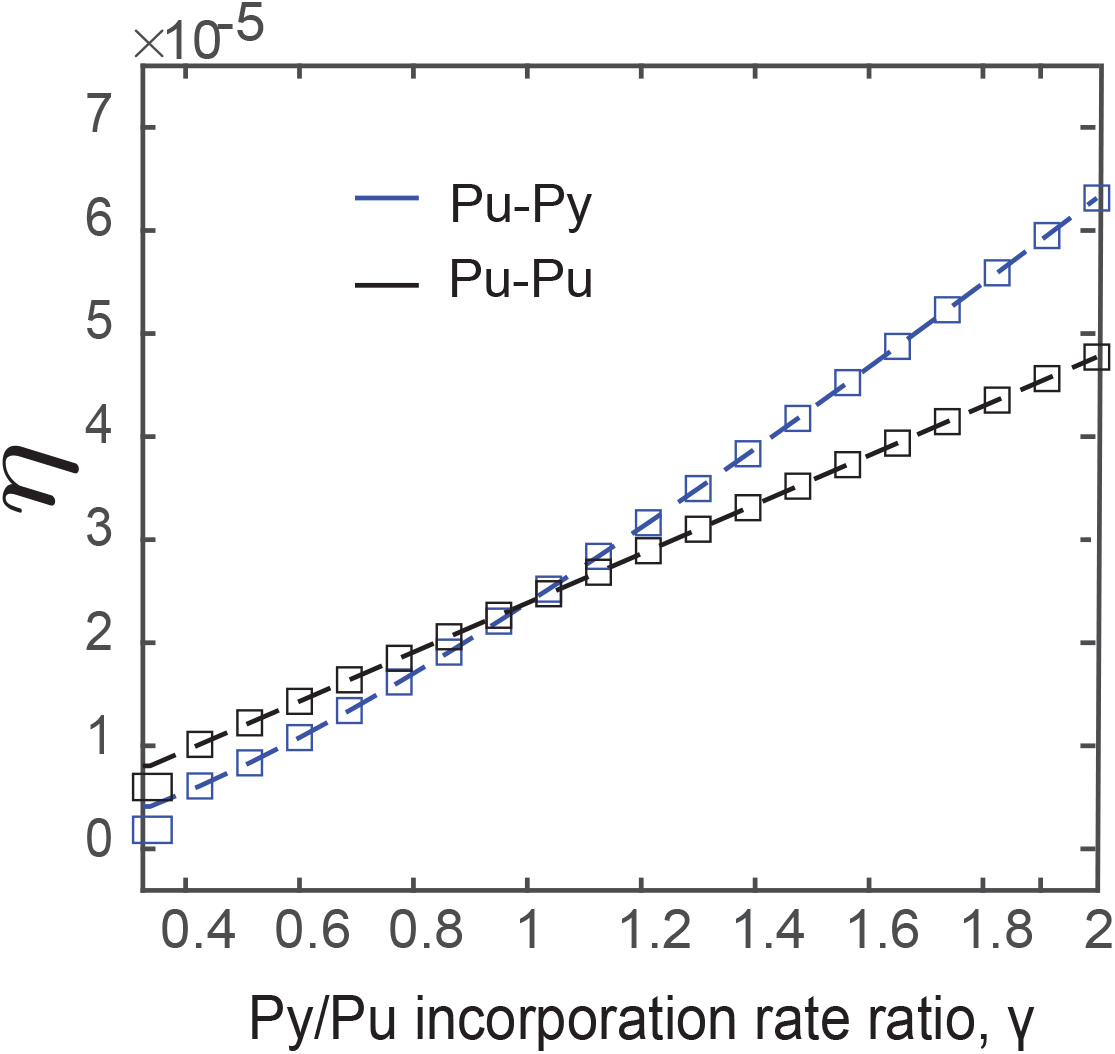
Comparison of numerical error from Eq. (4), methods (main manuscript) with the approximate analytical error rate from Eq. (S4) for two dimers: Pu-Pu, Pu-Py at the current and downstream positions of mRNA varying as a function of Py to Pu incorporation rate ratio, *γ*. Symbols and dashed lines respectively denote the numerical and approximate error result.

**Fig. S3.**
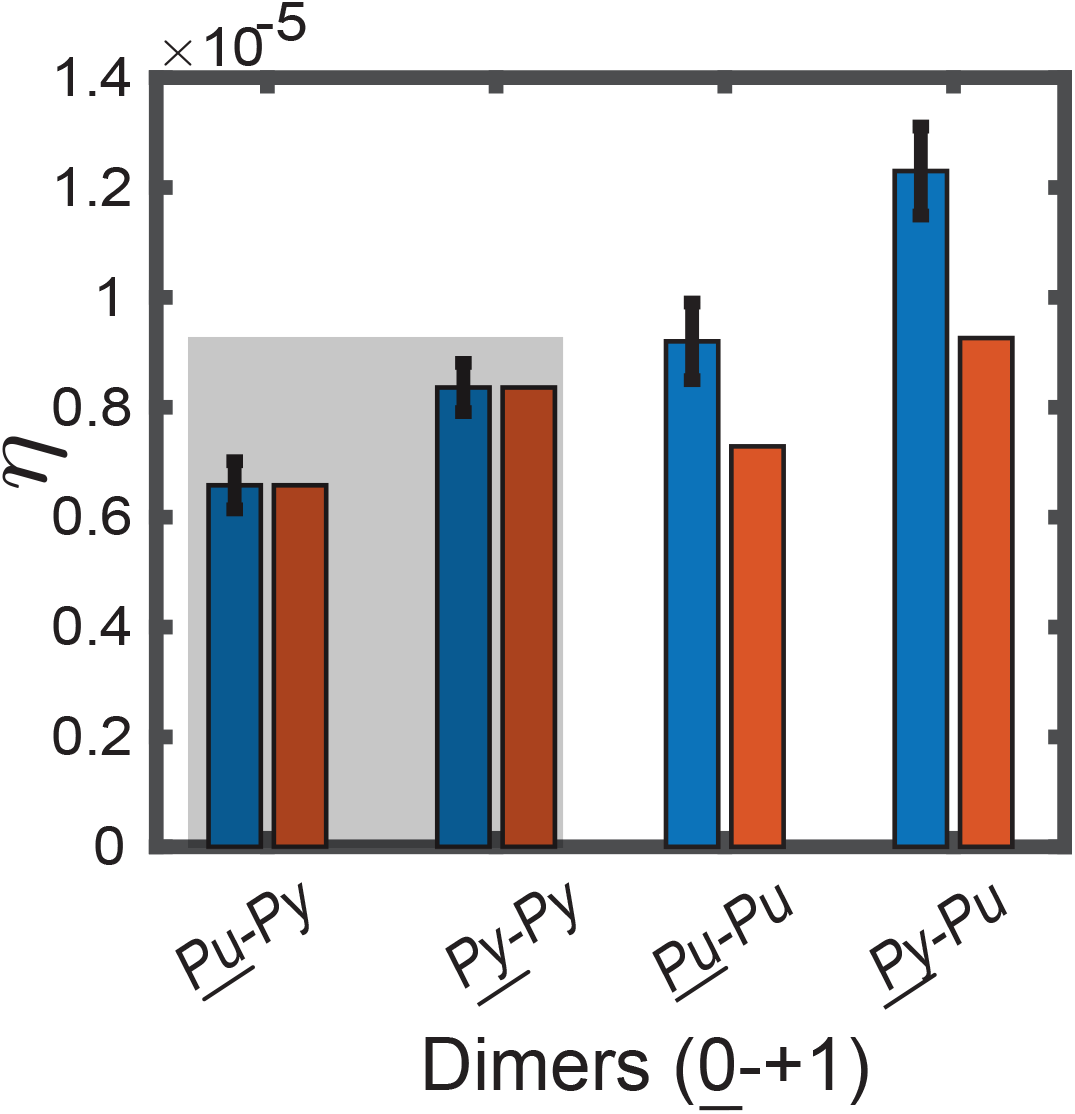
Fitting (shaded region) of theoretical errors (red bars) from the model with first-order neighbor effects and equal cleavage rates for purine and pyrimidines to the experimental data (blue bars). Theoretical errors (red bars) for the unfitted (unshaded region) dimers not aligned with the corresponding experimental data (blue bars). Error bars denote the standard error mean for the experimental data for the six samples (4).

**Fig. S4.**
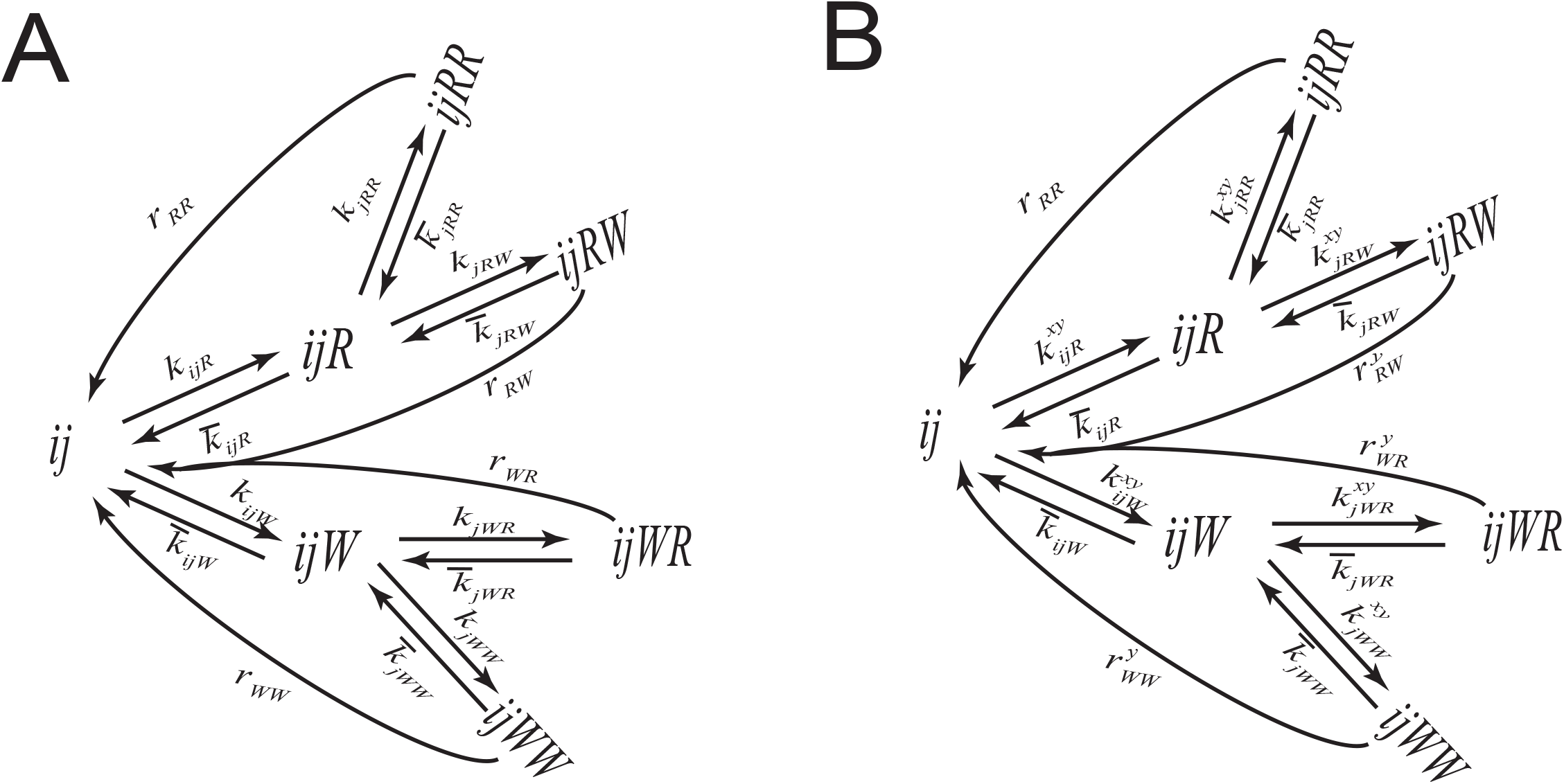
Schematic diagram of the two-element second-order biophysical model for transcription elongation process with dinucleotide cleavage and (A) without template-dependent kinetics (B) with template-dependent kinetics, where *ij* denotes the starting monomer. In 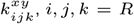, *W*, *x* denotes the nature, i.e., Pu or Py, of the previous incorporated nucleotide *j*, and *y* denotes the nature, i.e., Pu or Py, of the current incorporating nucleotide *k*. For no upstream effects, the rate 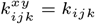, if *y* = Pu, and 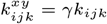, if *y* = Py. For the upstream effects, the rate 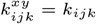, if *y* = Pu, *x* = Pu, 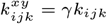, if *y* = Py, *x* = Pu, 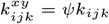, if *y* = Pu, *x* = Py, 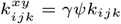, if *y* = Py, *x* = Py.

**Fig. S5.**
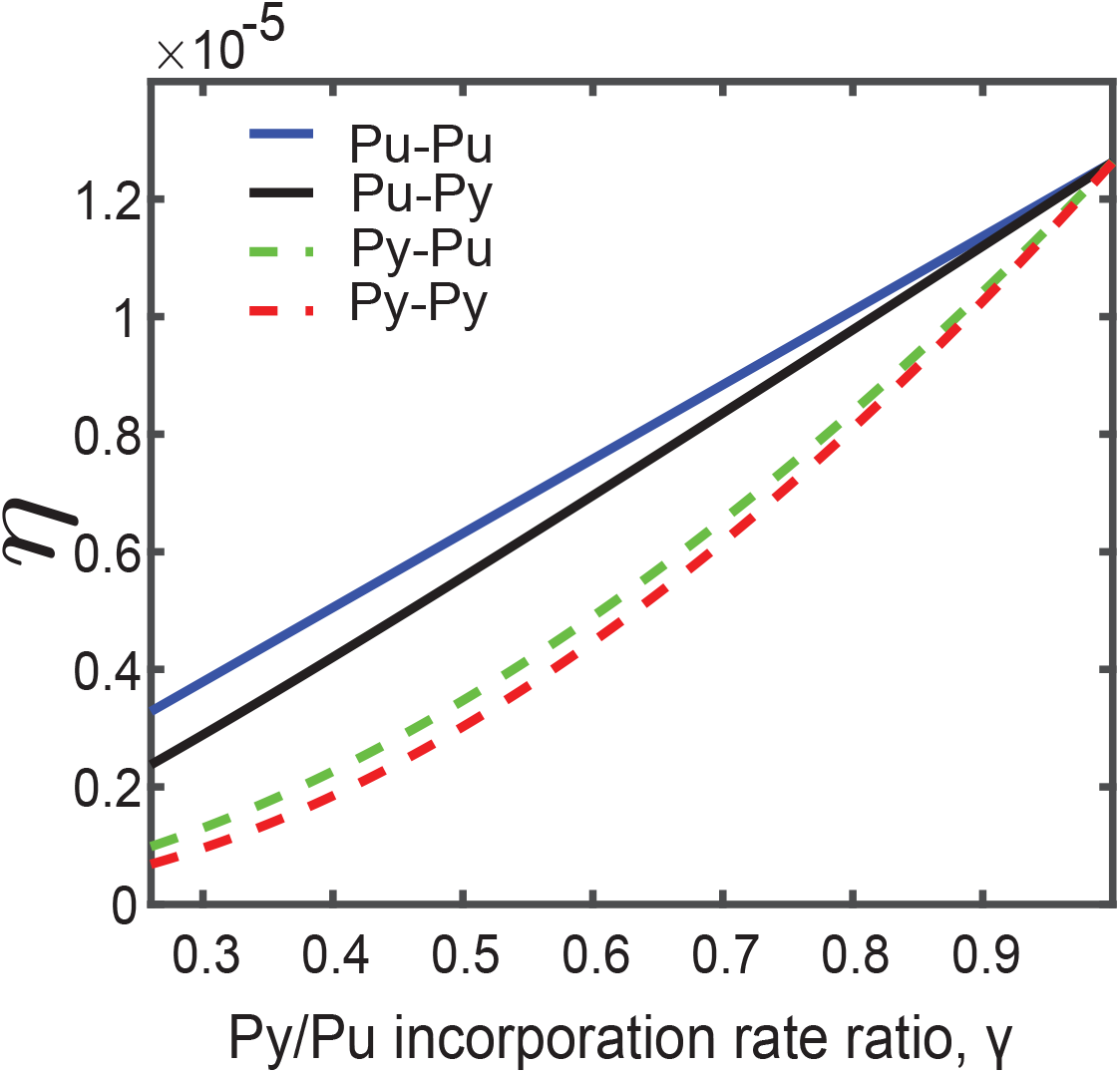
Variation of error from the two-element advanced model with a Pu at position 0 and four possible dimers occupying positions +1 and +2 on mRNA as a function of the incorporation factor, *γ*.

**Fig. S6.**
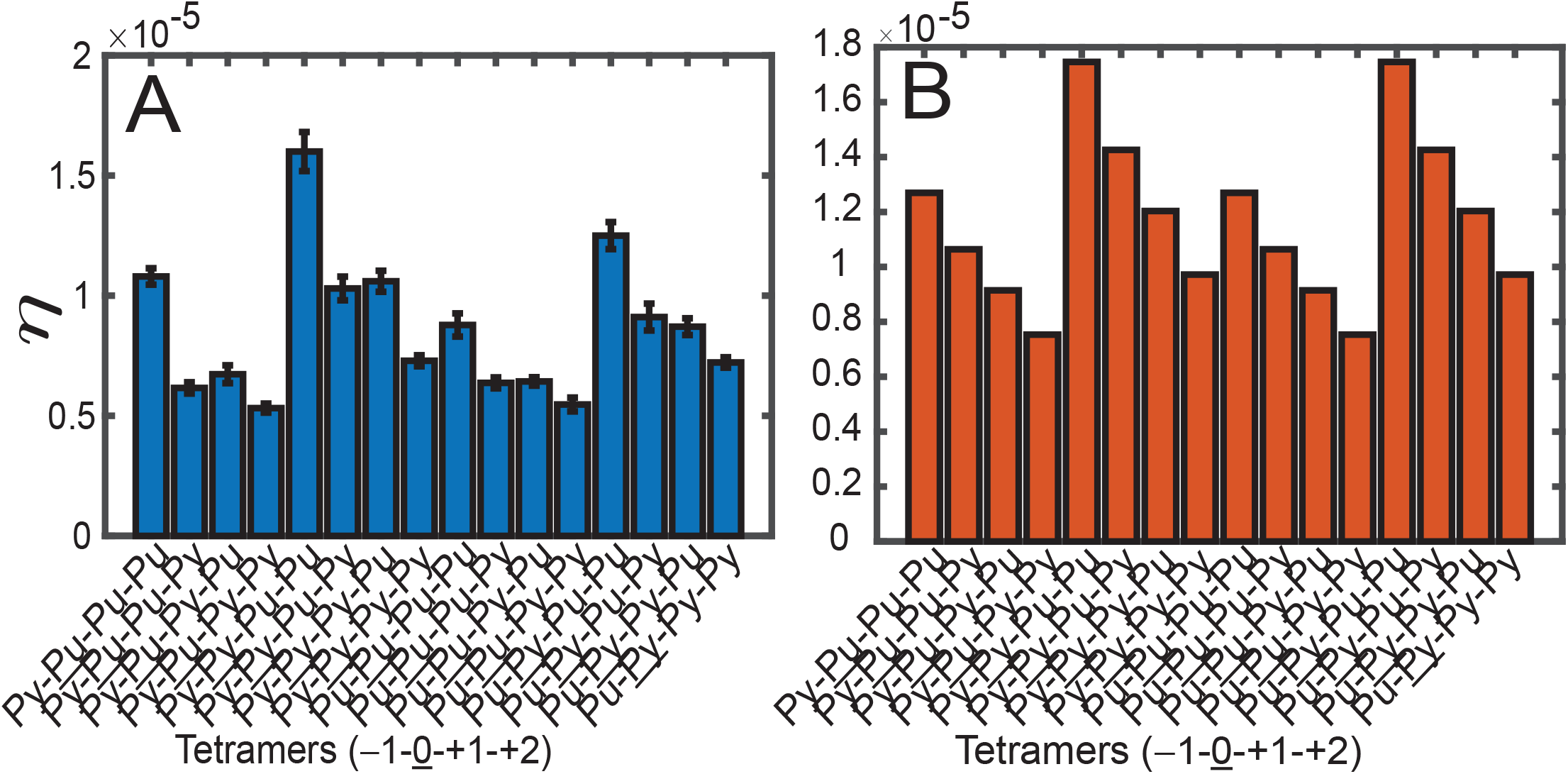
(A) Experimental error rates at a current position 0 depending on either purine or pyrimidine at positions -1, 0, +1, +2, obtained from the bioinformatics analysis of experimental data from Ref.(4). Error bars represent the SEM. (B) Theoretical error rates 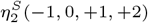 at a current position 0 depending on either purine or pyrimidine at positions -1, 0, +1, +2, obtained from the advanced model with second-order neighbor effects, two elements and no upstream nucleotide identity effect on the incorporation rates.

**Fig. S7.**
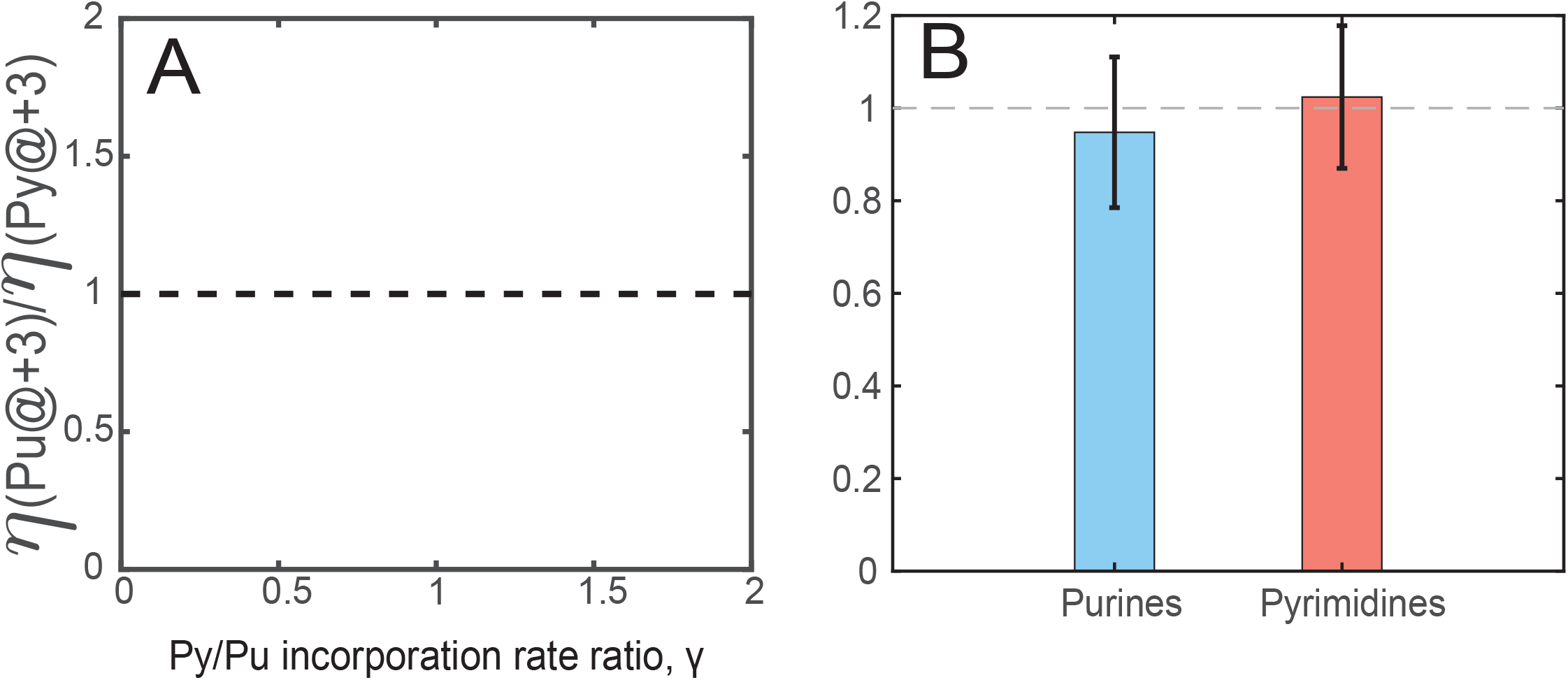
(A) Variation in the ratio of the theoretical error rate 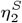 with Pu at +3 position to that of Py at +3 position *η*(Pu@ + 3)*/*(*η*_Py_@ + 3) with either Pu or Py at the current position, 0. (B) Variation in the ratio of the experimental error rate with Pu at +3 position to that of Py at +3 position.

**Fig. S8.**
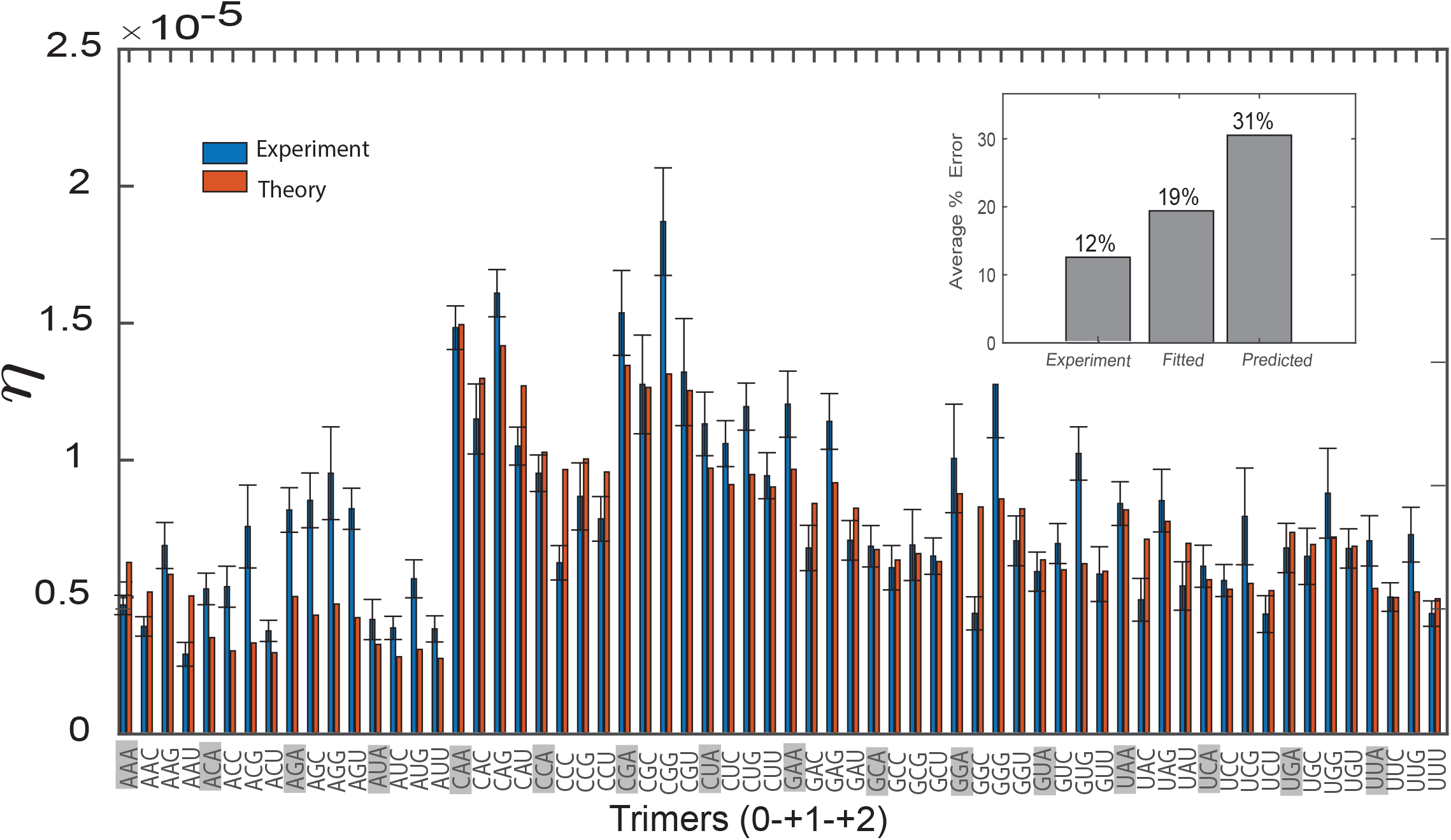
Fitting of theoretical error rates 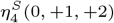 for the second-order biophysical model with four elements to the corresponding experimental data with cleavage rates depending on the nature of misincorporated nucleotide. Inset shows the average percentage of error for fitted, experimental, and unfitted data.

**Fig. S9.**
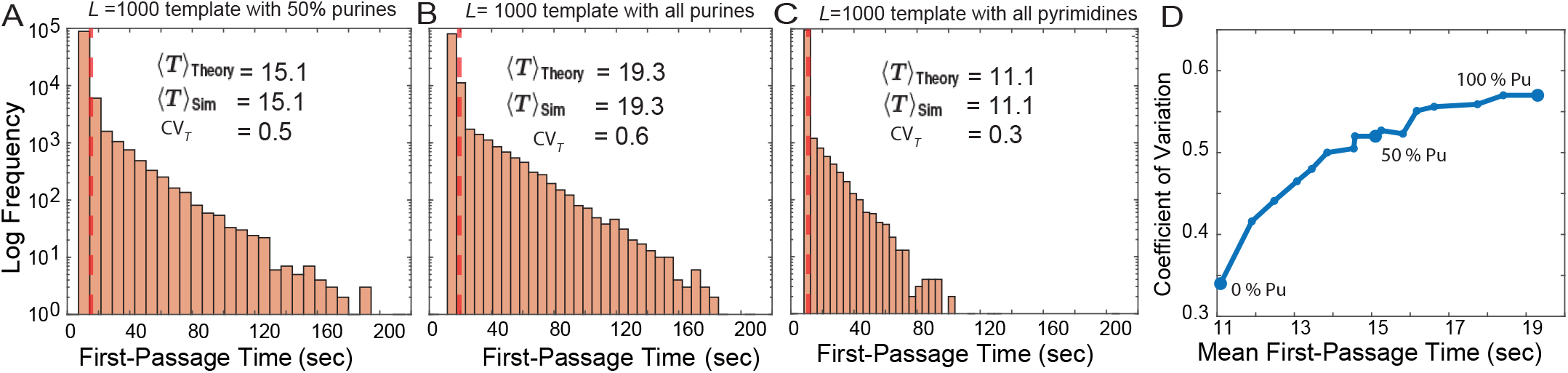
First-passage time (FPT) distributions for simulated transcription trajectories for a DNA template of length *L* = 1000 with varying base composition: (A) Random template with 50 % purines and % pyrimidines, (B) Purines-only template, (C) Pyrimidines-only template. For each case, the histogram shows the distribution of first-passage times from stochastic simulations (log-frequency scale), along with the corresponding mean first-passage time in the unit of seconds (sec) ⟨*T*⟩_Theory_ obtained from theory and ⟨*T*⟩_Sim_ obtained from simulations. The coefficient of variation (CV_*T*_) is also indicated, highlighting the influence of template composition on timing variability. (D) Mean first-passage time (MFPT) and coefficient of variation (CV_*T*_) as a function of purine content in the DNA template. Each point corresponds to a random template sequence with an increasing fraction of purines. The positive trend indicates that purine-rich templates are associated with both slower and more variable transcription.

**Table S1.** Kinetic parameters for the first-order biophysical model with two elements: Pu and Py.

| Parameter | Value $s^{-1}$ | Whether fitted or fixed | Role |
| --- | --- | --- | --- |
| $k_{RR}^{Pu}$ | 100 | Fitted | Correct incorporation rate for purine after $R$ at previous position |
| $k_{RR}^{Py}$ | $\gamma \times 100$ | Fitted $\gamma$ | Correct incorporation rate for pyrimidine after $R$ at previous position |
| $k_{RW}^{Pu}$ | .007 | Fixed | Misincorporation rate for purine after $R$ at previous position |
| $k_{RW}^{Py}$ | $\gamma \times .007$ | Fixed | Misincorporation rate for pyrimidine after $R$ at previous position |
| $k_{WR}^{Pu}$ | .01 | Fixed | Correct incorporation rate for purine after a misincorporation |
| $k_{WR}^{Py}$ | $\gamma \times .01$ | Fixed | Correct incorporation rate for pyrimidine after a misincorporation |
| $k_{WW}^{Pu}$ | $10^{-6}$ | Fixed | Misincorporation rate for purine after a misincorporation |
| $k_{WW}^{Py}$ | $\gamma \times 10^{-6}$ | Fixed | Misincorporation rate for pyrimidine after a misincorporation |
| $\bar{k}_{RR}$ | 5 | Fixed | Removal rate for a correct monomer after a $R$ monomer |
| $\bar{k}_{RW}$ | $9.65 \times 10^{-6}$ | Fixed | Removal rate for a wrong monomer after a $R$ monomer |
| $\bar{k}_{WR}$ | .00105 | Fixed | Removal rate for a correct monomer after a $W$ monomer |
| $\bar{k}_{WW}$ | $10^{-7}$ | Fixed | Removal rate for a wrong monomer after a $W$ monomer |
| $r_{RR}$ | .001 | Fixed | dinucleotide cleavage rate for correct dimer $RR$ |
| $r_{RW}^{Pu}, r_{WR}^{Pu}, r_{WW}^{Pu}, r_{Pu}$ | .038 | Fixed | Dinucleotide cleavage rate for wrong dimers with purines |
| $r_{RW}^{Py}, r_{WR}^{Py}, r_{WW}^{Py}, r_{Py}$ | $\phi \times .038$ | Fitted $\phi$ | Dinucleotide cleavage rate for wrong dimers with pyrimidines |

**Table S2.** Kinetic parameters for the advanced model with second-order neighbor effects and with two elements. For the upstream effect, the incorporation rate 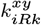 gets multiplied by the factor *ψ*, when previously incorporated nucleotide *x* is Py.

| Parameter | Value $s^{-1}$ | Fitted or Fixed | Role |
| --- | --- | --- | --- |
| $k_{RRR}^{xPu}$ | 100 | Fitted | Correct incorporation rate for purines after $RR$ |
| $k_{RRR}^{xPy}$ | $\gamma \times 100$ | Fitted $\gamma$ | Correct incorporation rate for pyrimidines after $RR$ |
| $k_{RRW}^{xPu}$ | .007 | Fixed | Misincorporation rate for purines after $RR$ |
| $k_{RRW}^{xPy}$ | $\gamma \times .007$ | Fixed | Misincorporation rate for pyrimidine after $RR$ |
| $k_{RW R}^{xPu}$ | 0.01 | Fixed | Correct incorporation rate for purines after $RW$ |
| $k_{RW R}^{xPy}$ | $\gamma \times 0.01$ | Fixed | Correct incorporation rate for pyrimidines after $RW$ |
| $k_{WRR}^{xPu}$ | 0.2 | Fixed | Correct incorporation rate for purines after $WR$ |
| $k_{WRR}^{xPy}$ | $\gamma \times 0.2$ | Fixed | Correct incorporation rate for pyrimidines after $WR$ |
| $\bar{k}_{RRR}$ | 5 | Fixed | Removal rate for any $R = \{Pu, Py\}$ monomer after $RR$ |
| $\bar{k}_{RRW}$ | $9.65 \times 10^{-6}$ | Fixed | Removal rate for any $W = \{Pu, Py\}$ monomer after $RR$ |
| $\bar{k}_{RW R}$ | .00105 | Fixed | Removal rate for any $R = \{Pu, Py\}$ monomer after $RW$ |
| $\bar{k}_{WRR}$ | .000105 | Fixed | Removal rate for any $R = \{Pu, Py\}$ monomer after $WR$ |
| $r_{RR}$ | .001 | Fixed | Dinucleotide cleavage rate for correct dimer $RR$ |
| $r_{RW}^{Pu}, r_{WR}^{Pu}, r_{WW}^{Pu}$ | .038 | Fixed | Dinucleotide cleavage rate for wrong dimers with purines |
| $r_{RW}^{Py}, r_{WR}^{Py}, r_{WW}^{Py}$ | .038 | Fitted | Dinucleotide cleavage rate for wrong dimers with pyrimidines |

**Table S3.**
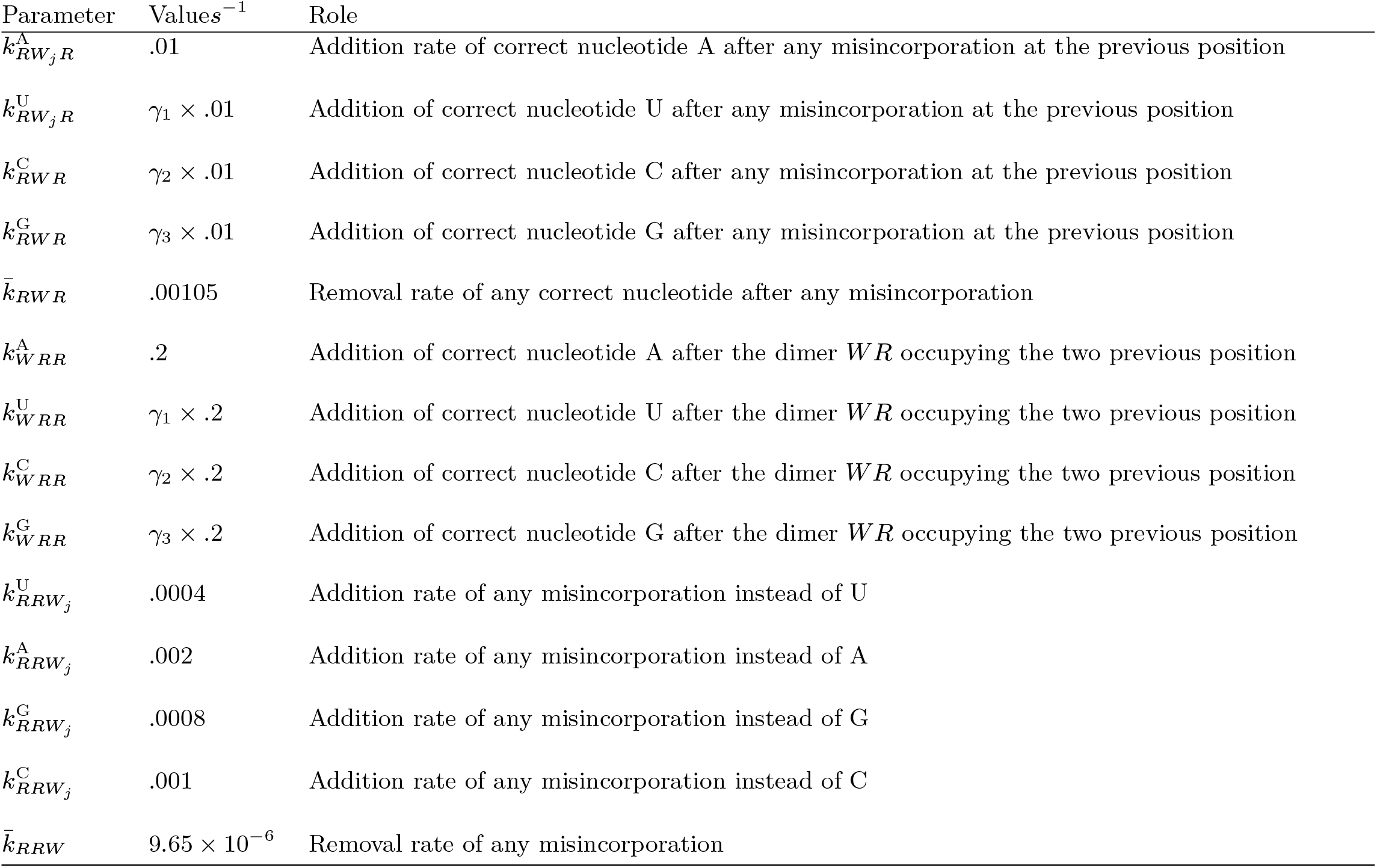
Kinetic parameters fixed for the computation of approximate theoretical error for the theoretical model with second-order neighbor effects and with four elements.

**Table S4.** Optimal kinetic parameters for the theoretical model with second-order neighbor effects and four elements obtained from fitting experimental data from Ref. (4).

| Parameter | Value | CV | Role |
| --- | --- | --- | --- |
| $k_{RRR}^A$ | $94.06\text{ s}^{-1}$ | 0.09 | Incorporation rate of A on mRNA given T on the DNA template |
| $\gamma_1$ | 0.64 | 0.02 | Incorporation rate ratio for U:A |
| $\gamma_2$ | 0.65 | 0.05 | Incorporation rate ratio for C:A |
| $\gamma_3$ | 0.99 | 0.03 | Incorporation rate ratio for G:A |
| $r_A$ | $.06\text{ s}^{-1}$ | 0.06 | Dinucleotide cleavage rate of misincorporations instead of correct A |
| $r_U$ | $.011\text{ s}^{-1}$ | 0.07 | Dinucleotide cleavage rate of misincorporations instead of correct U |
| $r_C$ | $.038\text{ s}^{-1}$ | 0.1 | Dinucleotide cleavage rate of misincorporations instead of correct C |
| $r_G$ | $.012\text{ s}^{-1}$ | 0.09 | Dinucleotide cleavage rate of misincorporations instead of correct G |

**Table S5.**
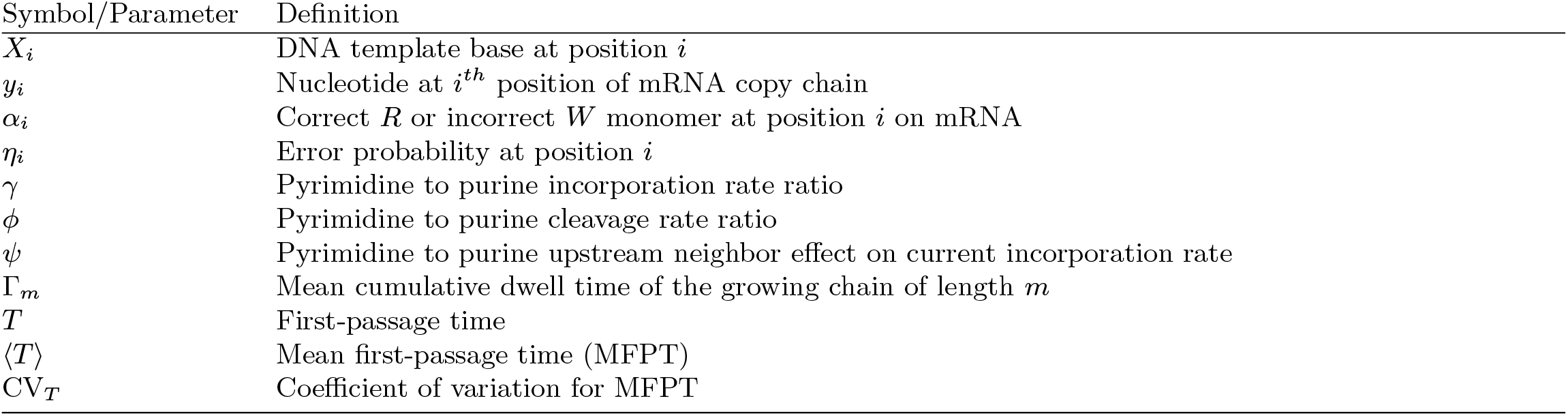
Definitions of some symbols and parameters.

| Symbol/Parameter | Definition |
| --- | --- |
| $X_i$ | DNA template base at position $i$ |
| $y_i$ | Nucleotide at $i^{th}$ position of mRNA copy chain |
| $\alpha_i$ | Correct $R$ or incorrect $W$ monomer at position $i$ on mRNA |
| $\eta_i$ | Error probability at position $i$ |
| $\gamma$ | Pyrimidine to purine incorporation rate ratio |
| $\phi$ | Pyrimidine to purine cleavage rate ratio |
| $\psi$ | Pyrimidine to purine upstream neighbor effect on current incorporation rate |
| $\Gamma_m$ | Mean cumulative dwell time of the growing chain of length $m$ |
| $T$ | First-passage time |
| $\langle T \rangle$ | Mean first-passage time (MFPT) |
| $CV_T$ | Coefficient of variation for MFPT |

